# Regenerative agriculture effects on biomass, drought resilience and ^14^C-photosynthate allocation in wheat drilled into ley compared to disc or ploughed arable soil

**DOI:** 10.1101/2025.09.04.674292

**Authors:** Nichola Austen, Elizabeth Short, Stefanie Tille, Irene Johnson, Richard Summers, Duncan D. Cameron, Jonathan R. Leake

## Abstract

Regenerative agriculture practices including leys and no-tillage facilitate biological reassembly of soil aggregates, increasing water, carbon and nutrient storage, but how this effects crop biomass, photosynthate partitioning, and drought resilience is unclear. To address this, we took monoliths growing semi-dwarf and taller wheat genotypes from 3-year plots that were ploughed or disc cultivated, or direct-drilled into a grass-clover ley and applied 35 kg N ha^-1^. Half the monoliths received a spring drought, then all were watered and the wheat ^14^CO_2_ pulse-labelled at stem elongation. The ley soil had lower bulk density, stored more water, and the taller wheat genotype maintained 8-fold higher proportion of water-stable macroaggregates despite unexpectedly having the smaller root biomass. Yields on the ley soil (3.74 t ha^-1^) were 77% −123% higher than on ploughed and disc cultivated soils, and unaffected by genotype or drought, despite its >70% reduction in root biomass. Of the ^14^C initially retained in wheat, 72% was in shoots, with root allocation decreasing by 75-90% in droughted ley soil, and at harvest soil retained <1% of the ^14^C, with significantly lower values for taller wheat, and ley. We conclude that soil health regeneration in the ley enhanced wheat yields, but reduced photosynthate allocation to root biomass under drought. Although taller wheat maintained better macroaggregation in ley soil, this was not explained by root biomass or photosynthate allocation and unexpectedly failed to increase soil ^14^C sequestration. We find no evidence that ley regeneration of macroaggregation enhances soil C sequestration under wheat, despite higher yields.

## 1. Introduction

Conventional production of high-yielding cereals grown intensively using ploughing harrowing and high fertilizer inputs is associated with deterioration of soil quality and functioning (Evans *et al*., 2020; Graves *et al*., 2015; Rickson *et al*., 2015). This results from depletion of soil organic matter, and physical disruption to water-stable soil macroaggregates that maintain macropore spaces through which air and water can rapidly move, at the same time storing organic matter and protecting it from microbial oxidation (Guest *et al*., 2022).

Wheat (*Triticum aestivum*) is the second most important cereal crop globally, after maize, with nearly 800 Mt produced annually and is an important food for both humans and livestock (Poutanen *et al*., 2021). In the UK wheat is the most widely grown crop, occupying about 38% of the cropped area in 2023 (Defra, 2024). Wheat breeding has focussed on yield enhancement, and modern varieties have the potential, under optimal soil, fertilizer, water and climate conditions, to deliver grain yields over 16 t ha^-1^, but the UK average is normally only about half this potential (Defra, 2023). One of the adverse consequences of the breeding of high-yielding modern wheat varieties is their small input of organic carbon via roots and root exudates, the forms of organic C that contribute a disproportionately large share of the organic matter that is stored in soils (Rasse *et al.,* 2005; Yang *et al.,* 2023). Wheat roots typically contribute about 0.4 t C ha^-1^ in a growing season (Sun *et al*., 2018), whereas perennial evergreen grass-clover leys can supply two and a half times more (∼1 t C ha^-1^; McNally *et al*., 2015). Since root-derived C typically has a residence time in soils 2.4 times that of shoots (Rasse *et al*., 2006), when wheat is frequently grown the limited C inputs may lead to deteriorating soil health (Kaloterakis *et al.,* 2024).

One of the crucial steps in the breeding of modern wheat varieties was the selection of dwarfing genes that reduced height and risk of lodging under the increasing ear weight achieved by selective breeding for enhanced yield (Velu *et al*., 2017). However, wheat dwarfing has been reported to associated with nearly 30% reductions in both root biomass, and shoot:root biomass ratios (Subira *et al.,* 2016), as well as reduced grain micronutrient concentrations (Velu *et al*., 2017). In addition, the breeding and selection of high yielding wheat varieties under conditions of abundant N and P fertilizer nutrient supply, and soil disturbance by ploughing and harrowing, may have selected against varieties that invest significant amounts of photosynthate into their mycorrhizal partners (Martín-Robles *et al*., 2017; Thirkell *et al*., 2022). Intensive arable cropping with ploughing and harrowing not only damages mycorrhizal hyphae, but also depletes earthworm populations (Edwards & Lofty, 1982), both of which are major contributors to soil aggregation and facilitate C storage in aggregates (Hallam & Hodson, 2020; Wilson *et al*., 2009). Intensive farming practices thereby both reduce the amounts of organic matter released into soils by roots and impair the functional biodiversity that assists in regenerating soil aggregate structures and sequestering organic C (Liu *et al.,* 2021; Or *et al.,* 2021; Zani *et al.,* 2021).

An increasing number of farmers are concerned about the escalating costs and unreliable returns from conventional intensive arable cropping, and the problems caused by soil degradation (Beacham *et al.,* 2023). Many in the UK are now adopting practices focussed on reducing input costs and regenerating soil health using practices that align with the broad concept of regenerative agriculture (Giller *et al*., 2021; Jaworski *et al*., 2023a,b; O’Donoghue *et al*., 2022; Beacham et al., 2023; RASE, 2025). The core principles of regenerative agriculture are to minimize soil disturbance, maintain living roots all year, return crop residue to soils, diversify rotations, and integrate grazing livestock (Brown, 2018; Jaworski *et al*., 2023b). One management practice, the reintroduction of legume-rich grass leys into arable rotations, can deliver all five core principles of regenerative agriculture, or four of the five if the leys are mown rather than grazed.

Recent research based on five conventional intensive arable and four permanent grassland fields at Leeds University farm in the SoilBioHedge (Hallam *et al*., 2020) and MycoRhizaSoil projects (Austen *et al.,* 2022) investigated effects of introducing mown grass-clover leys in arable rotations on soil health, biota and functioning. The leys led to rapid recovery in soil water-stable macroaggregates and organic carbon storage in these aggregates (Guest *et al*., 2022), paralleling recovery in earthworm population numbers (Prendergast-Miller *et al*., 2021; Llanos *et al*., 2025). At the same time, the leys showed significant improvements in soil hydrological functioning, with macroporosity and saturated hydraulic conductivity increasing with the more abundant earthworm populations (Hallam *et al*., 2020).

These soil functional changes are important as the main arable areas of eastern England are experiencing increasingly protracted periods of soil saturation due to climate change, exacerbated by impeded drainage where soil structures are damaged. These are also experiencing increasing droughts and high temperatures during the main growing season (Clarke *et al*., 2021), as exemplified by the drought in the spring of 2025 which was the driest in England since 1893 (Environment Agency, 2025). Resilience to weather extremes was increased for wheat direct drilled in grass-clover leys compared to long-term arable rotations, in an outdoor study using intact monoliths taken from 19-month-old leys and permanent arable plots in four adjacent fields in the SoilBioHedge trial (Berdeni *et al*., 2021). Wheat grown under flooding and moderate drought showing large yield improvements when direct drilled into leys, compared to when grown in ploughed arable soil (Berdeni *et al*., 2021). In contrast, in another European study on different soils, organic land management practices were not effective at alleviating drought yield losses compared to conventional agriculture (Sun *et al*., 2024), and a meta-analysis has suggested that regenerative practices do not consistently increase crop yields (Jordon *et al.,* 2022), although legume-rich leys were not considered in this, and crop yields were averaged over whole rotations including the ley years.

In MycoRhizaSoil the benefits of the leys on soil health for subsequent crops, including yield enhancement of 3.4 tonnes ha^-1^ was shown by a field trial comparing the performance of six wheat genotypes from the Avalon x Cadenza reference population, in which the parents and offspring lines differ in the extent of dwarfing (Bai *et al.,* 2013). The trial was established on long-term arable plots under ploughing and harrowing, three years of shallow disc cultivation, and direct drilling into a three-year grass-clover ley (Austen *et al*., 2022).

This study found in the wheat grown on the leys compared to permanent arable cropping with ploughing there were large increases in yield, 12 cm taller shoots, improved mycorrhizal colonization and reduced shoot pathology.

The effects of leys regenerating soil health (Berdeni *et al*., 2021; Hallam *et al*., 2020) and improving wheat growth in replicated field trials (Austen *et al*., 2022) raise important questions about the extent to which carbon flows from wheat into soil may be changed by reducing soil disturbance and regenerative field management practices. Of particular interest are the effects on wheat photosynthate allocation into soil from modern semi-dwarf rather than taller varieties and returning leys to arable cropping by direct drilling rather than ploughing, to reduce disruption of mycorrhizas and earthworms. We were also interested in how wheat dwarfing and leys and tillage management practices affected drought-resilience. Previous studies of C allocation from recent photosynthate in wheat using ^14^C or ^13^C labelling in pot, soil monolith, and field trials have been focused on conventional management systems in Australia (Atwell *et al*., 2002; Gregory & Atwell, 1991; Keith & Oades, 1986), and China (Sun *et al*. 2018; Sun *et al*., 2019). European studies on wheat using C-isotope pulse-labelling include comparisons of conventional with integrated (low N fertilizer use) management (Swinnen *et al*., 1995a,b) and crop rotation effects in conventionally managed cropping (Kaloterakis *et al*., 2024). These studies provide important insights into the growth-stage dependent root and shoot partitioning of photosynthate within wheat plants and into the rhizosphere and have shown significant effects of crop rotations (Kaloterakis *et al*., 2024).

However, they have left unaddressed the potential effects of regenerative farming practices such as direct drilling into fertility-building grass-clover leys on the partitioning of recent photosynthate in wheat of varieties that differ in extent of shoot dwarfing, and if these practices increase allocation of carbon into soil following drought.

We set out to test the following hypotheses: Reducing cultivation intensity from (a) annual inversion ploughing and harrowing to (b) three years of shallow disc cultivation and (c) direct drilling into a three-year grass-clover ley will:

1. Result in greater resilience to drought stress in the tall wheat than semi-dwarfed wheat grown on the ley as a result of having larger roots and better supporting soil structure and functioning, resulting in better yields.
2. Result in progressively increased organic C flux from wheat roots into soil C stores by the end of the growing season, due to reduced soil bulk density and improved soil macroaggregation, with these benefits being greatest under the non-dwarf wheat.

## 2. Materials and Methods

### 2.1 Field experiment overview

The field trial was set up at The University of Leeds farm, near Tadcaster, UK. The farm operates as a commercial, mixed livestock and arable farm. The soil is a loamy, calcareous Calcaric Endoleptic Cambisol of the Aberford Series (Cranfield University, 2019), underlain by a shallow bedrock of dolomitic limestone, which is representative of about 1,125 km^2^ of lowland England and Wales that is mainly used for arable cropping. The field used in the trial is located at 53◦ 51’ 44” N; 1◦ 20’ 35”, on a shallow south-facing slope of less than 10°.

The field was divided into six 20 m wide parallel strips that each ran downslope and were subdivided into four blocks that ran across the slope gradient, as shown in Austen *et al*., (2022). The strip closest to the field edge was used for small plot trials, the second strip was ploughed, harrowed and sown with ryegrass in November 2014, and oversown with red and white clovers in March 2015, with the resulting grass-clover ley maintained by mowing and removal of clippings. The remaining two pairs of strips were ploughed and harrowed, or shallow disc cultivated.

### 2.2 Wheat genotype selection

The parents and four double haploid wheat lines (AxC 22, AxC 53, AxC 157, and AxC 169) from the Avalon and Cadenza UK reference population (Wheat Genetic Improvement Network) provided by RAGT Seeds were sown annually in October – November 2015 to 2017. Cadenza can grow over 1 m tall, whereas Avalon is a semi-dwarf that is significantly shorter reaching only about 75 cm (Austen *et al*., 2022). These wheat lines were sown in annually ploughed and harrowed, and shallow disc cultivated plots described above and by Austen *et al*., (2022).

Two of the wheat lines that differed in dwarfing (*p* <0.001) were selected for the present study with AxC 157 attaining a mean height of 109 (± 1.25 standard error (s.e.)) cm in 2018 after direct drilling into ley, compared to only 85 (± 2.04 s.e.) cm with AxC 169 (Austen *et al*., 2022). These two wheat lines also had the most consistent contrasted rates of mycorrhization of roots in the summer of the first year of the trial (June 2016), with AxC 169 having mean arbuscule abundances of 7.7 - 11.5% of root length colonized, rising to 13.5% - 21.2% in AxC 157, although these differences were not significant (*p* > 0.05).

### 2.3 Field experiment plots from which soil monoliths were taken

The two wheat lines AxC 157 and AxC 169 were sown in early November 2017 into the main field experiment in replicate plots in each of the four blocks running across slope in the ploughed and harrowed and shallow disc cultivated strips, shown schematically in Figure 1. The parallel 3-year grass-clover ley strip was sprayed off with glyphosate in late October 2017, and the wheat lines were direct-drilled using a Simtech Aitchison grassland regeneration drill in November (see Figure 1). Guard strips of Skyfall wheat were sown at the sides and ends of most of the field blocks to protect the individual plots from wind damage, with strips of the original ley preserved from herbicide treatment (Figure 1). The wheat was grown in the field without use fungicides in the present study, but ferric phosphate slug pellets were used to protect establishing seedlings (full details in Austen *et al.,* 2022).

**Figure 1.**
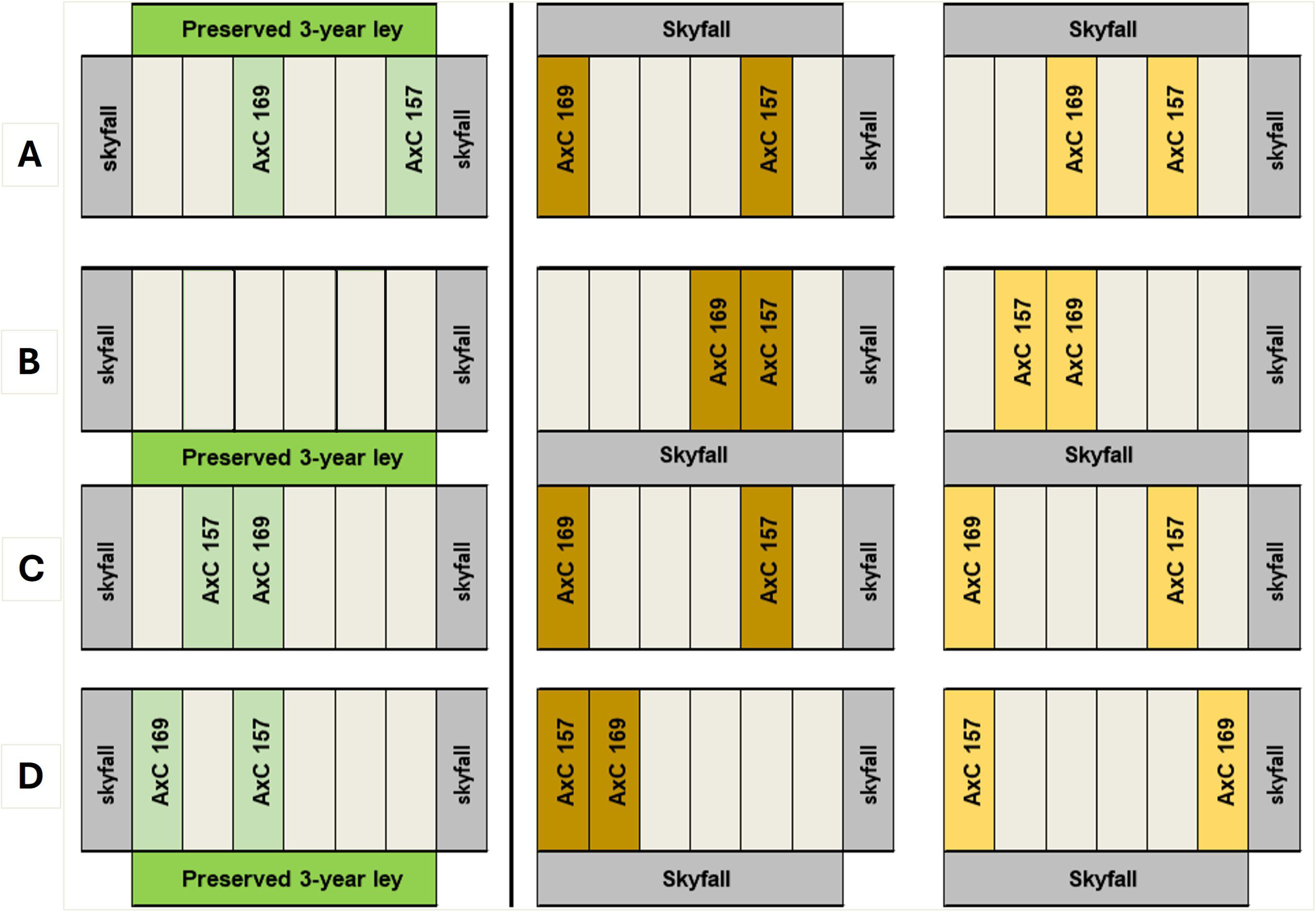
Schematic diagram of the design of the field trial with two wheat lines (AxC 157 and AxC 169) direct drilled into a 3-year grass-clover ley (green shading) or sown into three-year ploughed (brown shading) or disc cultivated (pale orange shading) wheat plots, protected by guard rows of Skyfall wheat (grey shading). From these plots soil monoliths were removed and transported to an outdoor compound at the University of Sheffield prior to ^14^ CO_2_ pulse-labelling of the wheat plants.

### 2.4 Soil monolith extraction from field site and their experimental treatments

In early April 2018, a total of 44 intact blocks of soil (monoliths) containing the established wheat plants were extracted from the field experiment (see Figure 1 for details the sampling locations, and supplemental info for photos of the monolith setup), following the methods used by Berdeni *et al.,* (2021). The monoliths fitted into 37 cm long, 27 cm wide, and 22 cm deep boxes with nine 10 mm drainage holes drilled into their bases and lined with 0.5 mm pore size nylon mesh to prevent loss of soil or earthworms. The monoliths were transported to the University of Sheffield and placed outdoors on drip-tray boxes on large trolleys insulated at the side using 10 cm thick Kingspan Thermawall board (see Supplementary Figure 1). The trolleys were placed in an open-sided polytunnel that provided a rain shelter as used by Berdeni *et al.,* (2021) to investigate wheat drought resilience in long-term arable ploughed versus direct drilled ley soil. Nitram ammonium nitrate fertilizer (34.5% N), matching that used in the field plots, was added in two equal splits in early April and in May immediately following watering to field capacity, to provide a total N equivalent to 35 kg N ha ^-1^, following Austen *et al.,* (2022).

The monoliths were initially watered twice weekly with 0.5 l of water (10 mm per week in addition to ambient rainfall) and were weighed using a digital hanging balance (precision ± 20 g), suspended below an engine hoist. From 13^th^ April 2018 for a period of 26 days, a rain shelter was installed over all of the monoliths, with half from each cultivation and wheat genotype treatment being assigned to a drought treatment and received no water, whilst the other half continued to receive 1 litre per week. For each wheat line there were 11 monoliths, with four replicates each for the ploughed and disc cultivated (one from each of blocks A-D), and three replicates for the former ley (from Blocks A, C, D). Immediately prior to imposing drought treatment (12^th^ April) all the monoliths were weighed, and then 19 days later, about three quarters of the duration of the drought, and three days (11^th^ May) after the rewetting of the monoliths to field capacity so the soil was unable to absorb any more water on 8^th^ May. The rain shelter was then removed. A final weight of the monoliths was recorded post ^14^C-labelling on 4^th^ June.

During the drought phase the soil moisture content of the top 7 cm of all the monoliths was assessed using a Delta-T time-domain reflectometry probe after 10 days, and then at a weekly interval for 2 weeks, ending on the day prior to rewetting on 8^th^ of May.

### 2.5 ^14^CO_2_ labelling of wheat in soil monoliths

Labelling of wheat was conducted over two consecutive days (14 - 15 May 2018), with 22 monoliths taken each day to a controlled environment greenhouse chamber with temperature of 20 °C. Each monolith was sealed in a large, clear polythene bag of circa. 80 L volume, using two pairs of bamboo canes inserted into the soil at the corners of the boxes and taped together at the top to form a tent-like structure over the plants which were in their stem elongation phase (AHDB Wheat Growth Guide, 2021), prior to ear formation (Supplementary Figure 2). A battery-operated small fan was attached to a wooden stake inserted vertically into the soil and positioned so the fan blades rotated in an air space close to the wheat shoots without touching them (see Supp. Fig. 2 for photographs). The fans were to ensure an air flow and homogenization of gas within the bags once sealed. The fans could be switched on and off through the bags (see Supp. Fig. 2 for photographs of the labelling set up). At the end of the monoliths closest to the greenhouse chamber central walkway a wooden stake was inserted vertically into the soil and a 2 ml Eppendorf tube containing 4 MBq of NaH^14^CO_2_ solution was firmly taped to the top of this. The lid of the tube could be carefully opened through the sealed bag, and 1.5 ml solution of 30% lactic acid added through the bag using a hypodermic needle attached to a 2 ml syringe, to liberate ^14^CO_2_. The needle hole in the bag was then immediately sealed using a pre-cut piece of waterproof tape.

The plants were incubated in the ^14^CO_2_ enriched atmosphere for three hours. Gas samples of 10 ml volume using a syringe fitted with a fine needle were taken immediately after liberating ^14^CO_2,_ at 30 minutes, 1 hour, 2 h and 3 h afterwards. At each sampling the syringes were pumped full and empty twice with gas from inside the labelling bag before the sample was taken, to ensure mixing of the air, and the needle holes in the labelling bags were then immediately sealed with waterproof tape. The gas samples were injected through a Subaseal into a scintillation vial with headspace gas evacuated, containing 10 ml of Carbosorb and 10 ml of Permafluor scintillation cocktail. At the end of the 3 h labelling periods the 22 monoliths were replaced outdoors into the open, the clear polythene bags cut off and removed, and the monoliths replaced on drip-trays on large trolleys and their sides surrounded by insulation board. This was repeated with the second set of 22 monoliths ^14^C labelled in the same way the following day.

The plants were maintained outdoors for 77 days after labelling (DAL), each monolith receiving 1 L of supplemental water per week, or 1.5 L if there was no rainfall. The plants continued to develop, flower, set seed and naturally senesced and ears ripened ready for harvest by August.

### 2.6 Plant shoot, root and soil samples taken for ^14^C quantification

Samples of 1-2 tillers out of a total of about 30 plants in each ^14^C labelled monolith were harvested two hours after the completion of the labelling, and again at three days after labelling (DAL), rapid oven dried at 80 °C for 48 h, cooled in a desiccator and weighed. At the three-day harvest an Eijelkamp bulk density corer was used to take 100 cm^3^ cores of 5.3 cm diameter from immediately under the sampled tillers sequentially from 0-7 and 8-15 cm depth, and the wheat roots were freed from soil, washed on a 1 mm sieve, blotted dry and fresh weight recorded before placing in a freezer. The frozen root samples were moved in batches to an Edwards freeze drier set to −50°C and −0.1 mbar vacuum, after which dry weights were taken.

The third and final harvest of the plant shoots, roots and soil samples took place in August (77 DAL). The plants were cut at the ground level, and the ears were separated from straw, and the grain and chaff were separated from the ears. The three components of the shoots were oven dried at 80 °C and then weighed.

The bulk density of whole monoliths (including stones) was determined from the measured depth of soil in each box multiplied by the internal length and breadth of the boxes and divided by the mass of soil in the box, considering the soil moisture content and mass of the box. The Eijelkamp bulk density corer was used to take 5.3 cm diameter cores from the centre of the monoliths from 0-7 cm depth, with soil fresh weight determined, then oven dried at 80 °C for 48 h, cooled in a desiccator and weighed to derive moisture content, and bulk density. An additional set of 0-7 cm soil core samples were taken from all the monoliths and wet-sieved into water-stable aggregate (WSA) fractions (>2000 µm, 1000-2000 µm, 250-1000 µm, 250-53 µm, and < 53 µm) following the method described by Guest *et al.,* (2022). The soil aggregate fractions were oven dried at 80 °C and weighed. Root biomass was estimated from soil cores taken sequentially at 0-7 and 7-15 cm depth of each monolith with roots extracted by teasing apart the soil and shaking soil from roots (which was retained and frozen), followed by the remaining roots and soil separated by washing on a 1 mm mesh and oven drying at 80 °C and weighing. The root free soil samples were later transferred in batches to a freeze dryer, as described earlier. These results were scaled to the whole monolith (0-15 cm) extrapolating from the core volumes.

### 2.7 Quantification of ^14^C in plant and soil samples

Plant tissue samples were ground to a homogenised powder by a ball mill (SPEX Cetriprep ball mill, Glen Creston Ltd.), and 30 mg sub-samples were weighed into a Combusto-Cone and burned in a sample oxidiser (Packard Sample Oxidiser, model 307). The evolved ^14^CO_2_ was trapped in 10 ml Carbosorb mixed with 10 ml Permafluor scintillation cocktail and radioactivity measured using a liquid scintillation analyser, Tri-Carb 3100TR, with a Quantum Smart programme. The following equations were used to determine the ^14^C recovered in the plant system where *eq.* 1 is the total amount of ^14^C from the sub sample of tillers harvested immediately after labelling and three DAL, scaled to the whole monolith and *eq*. 2 is the total amount of ^14^C recovered from the final biomass harvest.

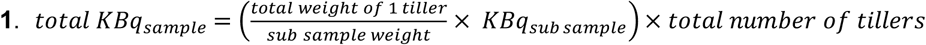

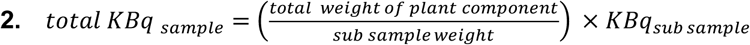

### 2.8 Statistical analyses

All statistical analyses were performed using MINITAB^®^ version 15 (Minitab, 2000) and Prism version 9.2.0 (GraphPad, 2021). Three-way ANOVA or two-way ANOVA was used to analyse the data with tillage type (direct drilled ley, disc cultivation, ploughed), water (ambient or drought), and wheat genotype (AxC 157, AxC 169) as the three factors. Both the independent and interactive effects of tillage type and wheat line were investigated. Where data did not meet the test requirements of homogeneity of variances across treatments, data was transformed by square root or natural log. In a few of the analyses of the soil aggregate sizes, there were a few extreme outlier values that were highlighted in the initial analysis, and these were removed for the final analysis. Tukey HSD post hoc tests were used to determine significant differences between the treatments (*p* <0.05).

## 3. Results

### 3.1 Effects of direct-drilled ley versus disc cultivation or ploughing on soil moisture content before, during and after drought treatment

The soil moisture content at 0-7 cm depth prior to the drought treatment showed no effect of wheat genotype, so we averaged across them (Fig. 2a). This showed that the soil moisture content in ley monoliths (36%) was significantly higher (*p* <0.05) than the disc cultivated or ploughed arable soil monoliths (31 and 30%, respectively), which did not differ significantly. With the imposition of drought, the soil moisture content fell to less than 10% over the first 10 days, the steepest fall occurring in the ley treatment, so that the three tillage treatments were no longer significantly different. In the monoliths with ambient watering the surface soil moisture content also fell to 18-22%, with the rank order of the treatments: direct-drilled ley, disc cultivated and ploughed being maintained but were no longer significantly different (*p* > 0.05). Over the following week the 0-7 cm soil moisture content stabilized and rose slightly in the ambient treatment, with no significant effects of the three tillage treatments, but continued to fall in the drought treatment, with the ley drying the most, but again the three tillage treatments did not differ significantly.

**Figure 2.**
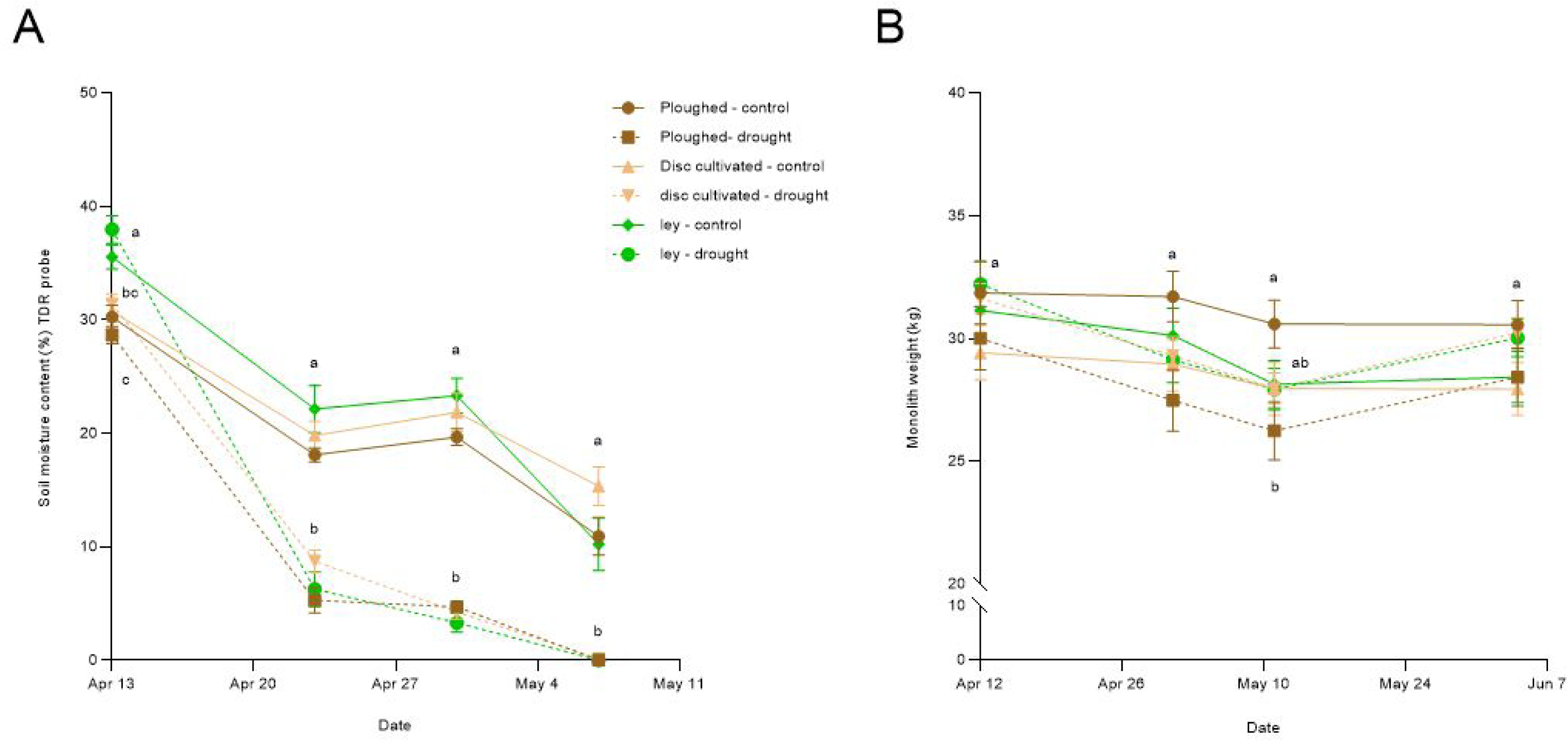
A) Effects of tillage treatments and drought (13 ^th^ April-8^th^ May) compared to watered controls on mean (± standard error) soil moisture content (% dry weight) of monolith 0-7 cm depth topsoil as measured by time domain reflectrometry (TDR) probe, averaging across the two wheat genotypes which did not differ significantly. Means sharing the same letter are not significantly different (*p* > 0.05) for comparisons on the same date (post hoc-Tukey test after 2-way ANOVA with tillage and water treatments as factors-see Table 2 for details). B) Effects of the tillage and drought treatments on whole soil monolith mean (± standard error) weight (kg) prior to drought treatment on 12^th^ April, during the drought, after rewetting to field capacity on 8^th^ May and after the ^14^C labelling on 4^th^ June, averaging across the wheat genotypes that did not differ significantly.

The weights of the monoliths were also recorded near the start of the study, during the drought treatment, and after rewetting to field capacity (Fig. 2b). As there were no effects of wheat genotypes these were again pooled to match Fig. 2a. Monolith weights prior to the drought treatment ranged by 10% from 32.22 kg in the ley to 29.43 kg in disc cultivated soils. The largest decrease in weight due to drought treatment was seen in the ploughed monoliths which had lost over 3.6 kg by the 11^th^ May, three days after rewetting to field capacity, and were significantly lighter (*p* = 0.048) by 15% than the watered ploughed controls at this time. Three days after rewetting, both disc cultivated (control and drought) and ley (control and drought) monoliths converged to weights of ∼ 28 kg (Fig. 2b & Table 1).

**Table 1.** Effects of tillage, genotype, and water treatment on monolith soil moisture, and monolith mass before during and after drought treatment analysed by two-way ANOVA. The results correspond to the data presented in Figure 2a and 2b. Bold text indicates significant value if *p* < 0.05.

| Figure and model | Date | Factor | *F*<br>(df) | *p* | Factor | *F*<br>(df) | *p* | Factor | *F*<br>(df) | *p* |
| --- | --- | --- | --- | --- | --- | --- | --- | --- | --- | --- |
| Fig 2a<br>2-way<br>ANOVA with<br>tillage and<br>water | 13.04.2018 | Tillage | 27.95<br>(2, 38) | **<0.001** | Water | 0.391<br>(1, 38) | *ns* | Tillage*<br>Water | 2.024<br>(2, 38) | *ns* |
|  | 23.04.2018 | Tillage | 2.941<br>(2, 38) | *ns* | Water | 167.7<br>(1, 38) | **<0.001** | Tillage*<br>Water | 1.748<br>(2, 38) | *ns* |
|  | 30.04.2018 | Tillage | 0.653<br>(2, 38) | *ns* | Water | 418.5<br>(1, 38) | **<0.001** | Tillage*<br>Water | 2.798<br>(2, 38) | *ns* |
|  | 07.05.2018 | Tillage | 2.430<br>(2, 38) | *ns* | Water | 127.2<br>(1, 38) | **<0.001** | Tillage*<br>Water | 2.196<br>(2, 38) | *ns* |
| Fig 2b<br>2-way<br>ANOVA with<br>tillage and<br>water | 12.04.2018 | Tillage | 0.51<br>(2, 38) | *ns* | Water | 0.27<br>(1, 38) | *ns* | Tillage*<br>Water | 1.96<br>(2, 38) | *ns* |
|  | 01.05.2018 | Tillage | 0.15<br>(2, 38) | *ns* | Water | 3.56<br>(1, 38) | *ns* | Tillage*<br>Water | 2.75<br>(2, 38) | *ns* |
|  | 11.05.2018 | Tillage | 0.13<br>(2, 38) | *ns* | Water | 3.47<br>(1, 38) | *ns* | Tillage*<br>Water | 3.284<br>(2, 38) | **0.0483** |
|  | 04.06.2018 | Tillage | 0.09<br>(2, 38) | *ns* | Water | 0.51<br>(1, 38) | *ns* | Tillage*<br>Water | 3.15<br>(2, 38) | *ns* |

**Table 2.** Effects of tillage, genotype, and water treatment on monolith soil bulk density after wheat harvest analysed by 3-way ANOVA. The results correspond to the data presented in Figure 3a and b. Bold text indicates significant value if *p* <0.05.

| Figure and model | Factor | *F*<br>(df) | *p* | Factor | *F*<br>(df) | *p* | Factor | *F*<br>(df) | *p* | Factor | *F*<br>(df) | *p* | Factor | *F*<br>(df) | *p* | Factor | *F*<br>(df) | *p* |  |  |
| --- | --- | --- | --- | --- | --- | --- | --- | --- | --- | --- | --- | --- | --- | --- | --- | --- | --- | --- | --- | --- |
| Fig. 3a<br>3-way ANOVA with tillage genotype and water | Tillage | 8.53<br>(2, 29) | <0.001 | Genotype | 0.53<br>(2, 29) | ns | Water | 6.02<br>(1, 29) | 0.02 | Tillage*<br>Genotype | 0.57<br>(2, 29) | ns | Tillage*<br>Water | 5.49<br>(2, 29) | 0.01 | Genotype*<br>Water | 0.01<br>(1, 29) | ns | Tillage*<br>Genotype*<br>Water | 3.36<br>(2, 29) |
| Fig. 3b<br>3-way ANOVA with tillage genotype and water | Tillage | 8.98<br>(2, 30) | <0.001 | Genotype | 0.00<br>(1, 30) | ns | Water | 1.85<br>(1, 30) | ns | Tillage*<br>Genotype | 1.49<br>(2, 30) | ns | Tillage*<br>Water | 4.38<br>(2, 30) | 0.021 | Genotype*<br>Water | 5.10<br>(1, 30) | 0.031 | Tillage*<br>Genotype*<br>Water | 0.58<br>(2, 30) |

Following the ^14^CO_2_ labelling, all droughted monoliths regained weight to almost match their mass prior to the drought, while the control monoliths that were watered throughout showed no significant change in weight from watering to field capacity on 8^th^ May (measured on 11^th^ May) and post labelling on 4^th^ June. Overall, there was a trend for the weight of the monoliths to decrease from 12^th^ April to 4^th^ June typically by 1-2 kg, despite the increase in biomass of the wheat that occurred during this time, indicating progressive drying of the soil.

### 3.2 Soil bulk density in direct drilled ley compared to disc cultivated and ploughed conventional arable soil

Whole monolith soil bulk density (BD) (Fig. 3a) measured after the wheat harvest only showed significant effects of tillage (*p* < 0.001), drought (*p* < 0.02), an interaction between tillage and drought (*p* = 0.01) and an interaction between wheat variety, tillage and drought (*p* < 0.05), see Table 2. Averaging across wheat varieties and watering treatments, the direct drilled ley (1.13 g cm^3^ ± 0.01 s.e.) was significantly less compacted than the disc and ploughed soils (both 1.23 g cm^3^ ±0.02 s.e.). Drought reduced the average BD from 1.27 g cm^3^ (±0.02 s.e.) to 1.18 g cm^3^ (± 0.02 s.e.) in the ploughed soils, and from 1.27 g cm^3^ (±0.02 s.e.) to 1.16 g cm^3^ (±0.02 s.e.) in disc cultivated soils. The interaction between tillage and drought was due to drought having no significant effect on the BD in the direct drilled ley samples (ambient 1.16 (± 0.01 s.e.) vs 1.18 g cm^3^ (± 0.02 s.e.) with drought The only effect of wheat variety was in the three-way interaction arising from AxC 157 in the direct drilled ley having slightly lower bulk density under ambient watering than in the ploughed or disc cultivated, but there were no differences under drought, whereas all other combinations of variety, watering and tillage treatments showed no significant differences (Fig. 3a).

**Figure 3.**
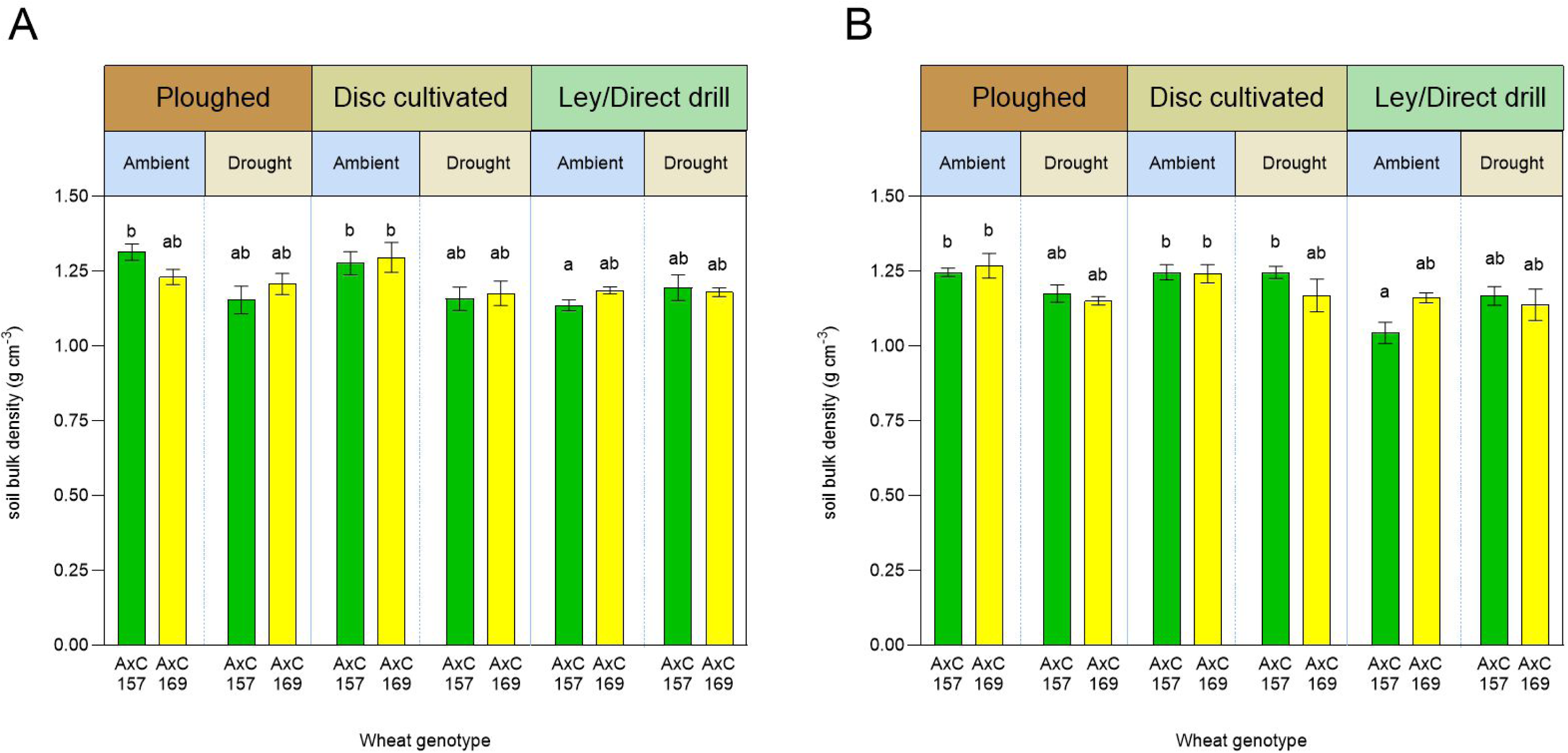
Soil bulk density (g cm^-3^) responses to tillage, drought and wheat variety in A) Whole monoliths, B) the 0-7 cm monolith topsoil. Error bars show ± 1 SEM. Means sharing the same letter are not significantly different (*p* > 0.05) in post hoc - Tukey test after 3-way ANOVA.

For the 0 – 7 cm topsoil (Fig. 3b), the BD was significantly affected by tillage treatment (*p* < 0.001), the interaction between wheat variety and water treatment (*p* = 0.031), and the interaction between tillage and water treatment (*p* = 0.021), the full ANOVA results being given in Table 2. As with the whole monoliths, the direct drilled ley had the lowest mean BD of 1.13 g cm^3^ (± 0.02 s.e.) and there was no significant difference between the ploughed (1.21 g cm^3^ ± 0.01 s.e.) and disc cultivated (1.23 g cm^3^ ± 0.01 s.e.) monoliths. The interaction between wheat variety and water treatment arose because AxC 157 showed no difference between ambient (1.19 g cm^3^ ± 0.03 s.e.) and drought (1.19 g cm^3^ ± 0.01 s.e.) treatments, whereas with AxC 169 the drought treatment reduced mean bulk density from 1.22 (± 0.02 s.e.) to 1.15 g cm^3^ (± 0.02 s.e.).

### 3.3. Differences in abundance of different sized water-stable aggregates in direct drilled ley compared to disc cultivated and ploughed conventional arable soil

Three-way ANOVA on square root transformed data (with one outlier removed) showed that the proportion of the 0-7 cm soil in WSA > 2 mm was significantly affected by tillage (*p* < 0.001), with the direct drilled ley supporting a higher proportion of soil in these large macroaggregates (15.9%) compared to disc cultivated (5.2%) and ploughed (4.2%), which were not significantly different (Fig. 4, see Table 3 for full ANOVA results). There was an interaction between wheat variety and tillage (*p* = 0.002) which was driven by AxC 157 grown on the direct drilled ley having over seven times more WSA > 2 mm than the disc or ploughed soils with the same wheat variety (Tukey test *p* < 0.001), whereas with AxC 169 the decrease in these macroaggregates from the direct drilled ley to the disc and ploughed soil were much smaller and not significant (Tukey test *p* > 0.05). A significant three-way interaction between tillage, watering treatment, and wheat variety (*p* = 0.003) also arose from the greater abundance of the > 2 mm WSA in the direct drilled ley with ambient watering growing AxC 157.

**Figure 4.**
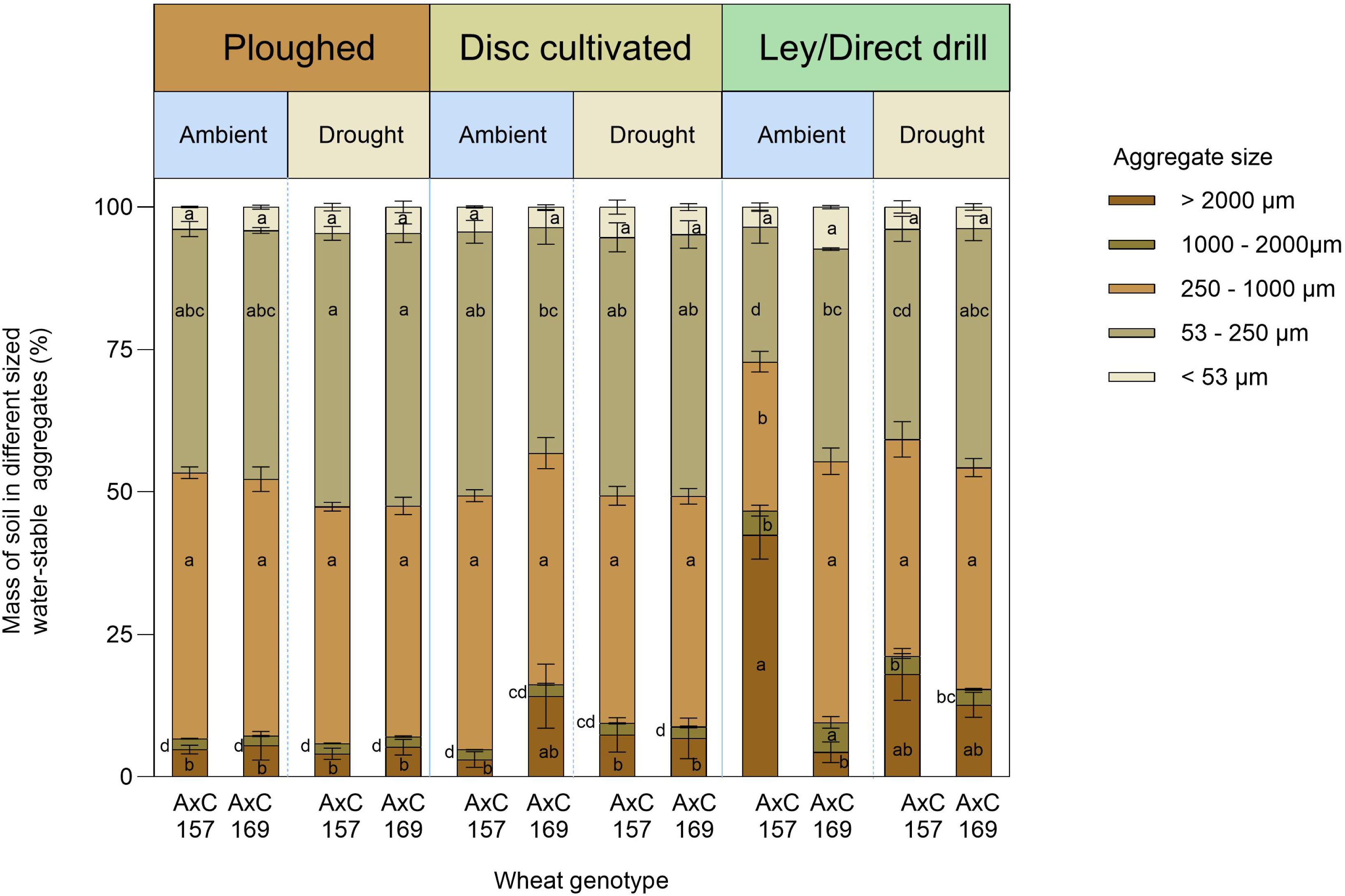
Water stable aggregate sizes of soil from monoliths used to grow AxC157 and AxC 169 with the different tillage and water treatments. Error bars show ± 1 SEM. Means sharing the same letter are not significantly different (*p* > 0.05) (post hoc-Tukey test after 3-way ANOVA with tillage, water treatment and wheat genotype as factors

**Table 3.** Effects of tillage, genotype, and water treatment on proportion of soil mass in different water-stable aggregate fractions after the wheat harvest, analysed by 3-way ANOVA, and corresponding to the mean values presented in Fig. 4.

| WSA* | Factor | F<br>(df) | p | Factor | F<br>(df) | p | Factor | F<br>(df) | p | Factor | F<br>(df) | p | Factor | F<br>(df) | p | Factor | F<br>(df) | p |
| --- | --- | --- | --- | --- | --- | --- | --- | --- | --- | --- | --- | --- | --- | --- | --- | --- | --- | --- |
| >2000µm | Tillage | 11.42<br>(2, 61) | <0.001 | Genotype | 2.83<br>(1, 61) | ns | Water | 0.95<br>(1, 61) | ns | Tillage*<br>Genotype | 7.24<br>(2, 61) | 0.002 | Tillage*<br>Water | 0.08<br>(2, 61) | ns | Genotype*<br>Water | 1.12<br>(1, 61) | ns |
| 1000-2000µm | Tillage | 96.31<br>(2, 59) | <0.001 | Genotype | 1.91<br>(1, 59) | ns | Water | 4.7<br>(1, 59) | 0.034 | Tillage*<br>Genotype | 1.32<br>(2, 59) | ns | Tillage*<br>Water | 7.38<br>(2, 59) | <0.001 | Genotype*<br>Water | 14.04<br>(2, 59) | <0.001 |
| 250-1000µm | Tillage | 7.97<br>(2, 61) | 0.001 | Genotype | 3.82<br>(1, 61) | ns | Water | 1.71<br>(1, 61) | ns | Tillage*<br>Genotype | 9.14<br>(2, 61) | <0.001 | Tillage*<br>Water | 2.76<br>(2, 61) | ns | Genotype*<br>Water | 3.45<br>(1, 61) | ns |
| 53-250µm | Tillage | 17.81<br>(2, 61) | <0.001 | Genotype | 2.22<br>(1, 61) | ns | Water | 13.59<br>(1, 61) | <0.001 | Tillage*<br>Genotype | 5.72<br>(2, 61) | 0.0053 | Tillage*<br>Water | 5.72<br>(2, 61) | ns | Genotype*<br>Water | 0.06<br>(1, 61) | ns |
| <53µm | Tillage | 0.12<br>(2, 61) | ns | Genotype | 0.75<br>(1, 61) | ns | Water | 0.00<br>(1, 61) | ns | Tillage*<br>Genotype | 1.81<br>(2, 61) | ns | Tillage*<br>Water | 2.21<br>(2, 61) | ns | Genotype*<br>Water | 1.54<br>(1, 61) | ns |

Bold text indicates significant value if $p < 0.05$ . \* = Water stable aggregate fraction

The 1-2 mm WSA were the least abundant of the aggregate fractions but still showed highly significant effects of tillage (*p* < 0.001) with the direct drilled ley supporting approximately double the abundance of these aggregates than the disc or ploughed treatments, which were not significantly different (*p* > 0.05). There was also a significant overall effect of drought treatment (*p* = 0.034) which reduced the abundance of these aggregates by 10%. A significant interaction was found between the effects of wheat variety and drought treatment, with no significant effect of drought in line AxC 157, (Tukey test *p* > 0.05) but a 21% decrease from drought with AxC 169 (Tukey test *p* < 0.001). There was also a highly significant interaction between the effects of tillage and drought treatments (*p* = 0.001), due to drought reducing the abundance of the 1-2 mm WSA by 25%, only in the direct drilled ley treatment. A three-way interaction between wheat variety, tillage and drought treatments was highly significant (*p* < 0.001), driven by the higher abundance of 1-2 mm WSA with AxC 169 in the direct drilled ley under ambient water supply.

The 0.25-1 mm WSA typically accounted for about 40% of the soil mass, except for the direct drilled ley with ambient water on which AxC 157 was grown, where it fell to 26% and was significantly different (Tukey test *p* < 0.05) to all other treatment combinations (Fig. 4). Overall, there was a significant effect of tillage (*p* < 0.001) because of direct drilling reducing the abundance of these aggregates to 37% of the soil mass compared to 41% and 43% in the disc cultivated and ploughed soils, respectively, which did not differ from each other (Tukey test *p* > 0.05).

The WSA fraction of 53-250 μm also typically accounted for about 40% of the soil mass and again showed significant effects of tillage *(p* < 0.001). The direct drilled ley showed only 34.9% of soil in this fraction, compared to 44% in the disc cultivated and 45.6% in the ploughed treatments, which were not significantly different to each other (*p* > 0.05). There was also a highly significant effect of drought (*p* <0.001) increasing the soil mass to 44.4% compared to 38.9% under ambient conditions. There was an interaction between wheat variety and tillage (*p* <0.005), arising from AxC 157 in the direct drilled ley having much lower abundance of this soil fraction (35%) than when grown in disc (45.9%) or ploughed (45.4%) soil, whereas with AxC 169 there were no significant effects of tillage.

The smallest fraction of the wet-sieved soil (< 53 μm) accounted for ∼ 5-7% of the total soil mass across all treatments. The results of the three-way ANOVA yielded no significant effects of tillage, wheat genotype or water treatment, nor any significant interactive effects of these variables (Fig. 4, Table 3; *p* > 0.05).

### 3.4 Wheat shoot biomass and grain yields in direct drilled ley compared with conventional disc cultivated and ploughed soil mesocosms

The ley and tillage methods significantly affected total above ground biomass (*p* < 0.001). Grain yields were only affected by tillage (*p* < 0.001), with yields on the ley of 3.74 t ha^-1^ (± 0.01 s.e.) compared to 2.11 t ha^-1^ (± 0.11 s.e.) in the ploughed plots and 1.67 t ha^-1^ (± 0.12 s.e.) in the disc cultivated plots, averaging across genotypes and water treatments which were not significant (Table 4, Supp. Table 2). These biomass differences due to ley and tillage were also seen in the chaff and straw (Fig. 5, Table 4, Supp. Table 2). Only the straw biomass of the two wheat lines was affected by the independent effects of genotype (*p* = 0.013) and water treatment (*p* = 0.001). AxC 169 had a lower straw biomass than AxC 157 (Fig. 5), with the droughted monoliths following the same trend. Averaging across wheat lines and tillage, the droughted monoliths gave 15% lower straw yield than the ambient monoliths (*p* = 0.001).

**Figure 5.**
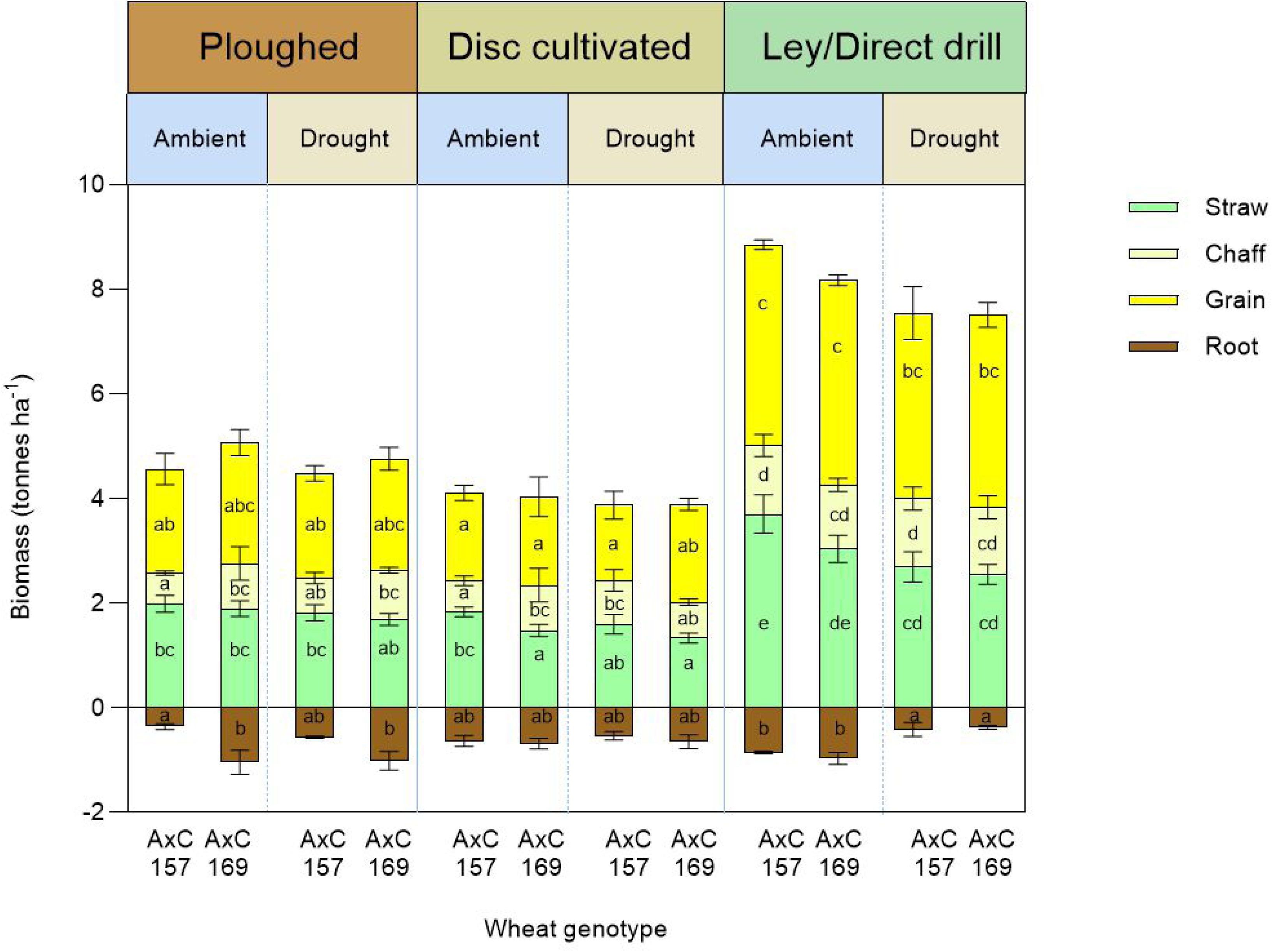
Biomass of wheat shoots (grain, chaff and straw) and roots by variety (AxC 157 vs AxC 169), tillage and drought treatment combinations. Error bars show ± 1 SEM. Means sharing the same letter within shoot components or roots are not significantly different (*p* > 0.05) in Tukey tests after three-way ANOVA.

**Table 4.** Effects of tillage, genotype, and water treatment on wheat shoot and root biomass after the wheat harvest, analysed by 3-way ANOVA, and corresponding to the mean values presented in Fig. 5. Bold text indicates significant value if *p* <0.05.

| Figure and model | Plant fraction | Factor | *F* (df) | *p* | Factor | *F* (df) | *p* | Factor | *F* (df) | *p* | Factor | *F* (df) | *p* | Factor | *F* (df) | *p* | Factor | *F* (df) | *p* |
| --- | --- | --- | --- | --- | --- | --- | --- | --- | --- | --- | --- | --- | --- | --- | --- | --- | --- | --- | --- |
| Fig. 5<br>3-way ANOVA with tillage genotype and water | Grain | Tillage | 66.13 (2, 32) | **<0.001** | Genotype | 1.50 (1, 32) | ns | Water | 0.71 (1, 32) | ns | Tillage* Genotype | 0.06 (2, 32) | ns | Tillage* Water | 0.22 (2, 32) | ns | Genotype* Water | 0.11 (1, 32) | ns |
|  | Chaff | Tillage | 8.98 (2, 32) | **0.001** | Genotype | 0.71 (1, 32) | ns | Water | 0.17 (1, 32) | ns | Tillage* Genotype | 0.76 (2, 32) | ns | Tillage* Water | 0.02 (2, 32) | ns | Genotype* Water | 0.31 (1, 32) | ns |
|  | Straw | Tillage | 64.86 (2, 32) | **<0.001** | Genotype | 6.87 (1, 32) | **0.013** | Water | 13.11 (1, 32) | ns | Tillage* Genotype | 0.68 (2, 32) | ns | Tillage* Water | 2.88 (2, 32) | ns | Genotype* Water | 0.87 (1, 32) | ns |
|  | Roots | Tillage | 0.95 (2, 29) | ns | Genotype | 8.52 (1, 29) | **0.007** | Water | 4.55 (1, 29) | **0.042** | Tillage* Genotype | 4.94 (2, 29) | **0.014** | Tillage* Water | 4.97 (2, 29) | **0.014** | Genotype* Water | 0.46 (1, 29) | ns |

### 3.5 Varietal, tillage and drought effects on wheat root biomass and root: shoot ratios at harvest

Averaging across tillage and water treatments, AxC 169 unexpectedly produced significantly more root biomass than AxC 157 (*p =* 0.007) (Fig. 5; Table 4). The varietal differences in root biomass were greatest in the ploughed treatments, and not different in the minimal tillage and ley plots giving a genotype by tillage treatment interaction (*p* < 0.05, Fig. 5, Table 4). Drought had no effect on root biomass in ploughed and minimal tillage plots, but reduced root biomass in the direct drilled ley by 70% in AxC 157 and and 83% in AxC 169 compared to ambient ley treatment in each case, giving a tillage by drought treatment interaction (*p* < 0.05; Fig. 5; Table 4).

There was an overall significant difference (*p* < 0.05; nat. log transformed data) between the root:shoot biomass ratios of the wheat varieties (Fig. 6; Table 5), but contrary to our original expectations, the semi-dwarf AxC 169 had a greater proportion of biomass in its roots (arithmetic mean 0.16 ± 0.02 s.e.) than the taller AxC 157, (0.11 ± 0.01 s.e.).

**Figure 6.**
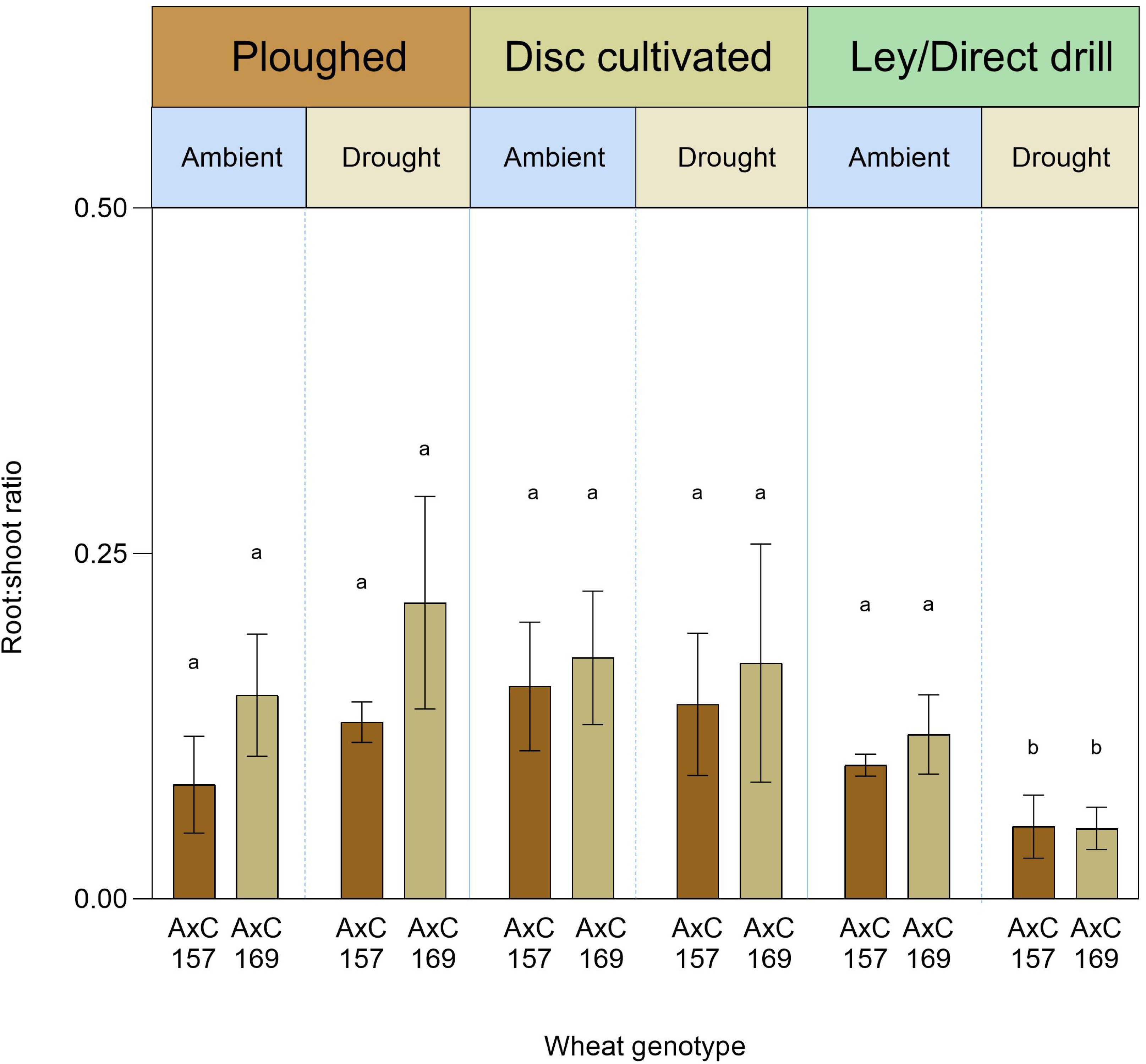
The mean root to shoot weight ratio (± standard error) of the two wheat genotypes in response to the three tillage and drought stress treatments.

**Table 5.**
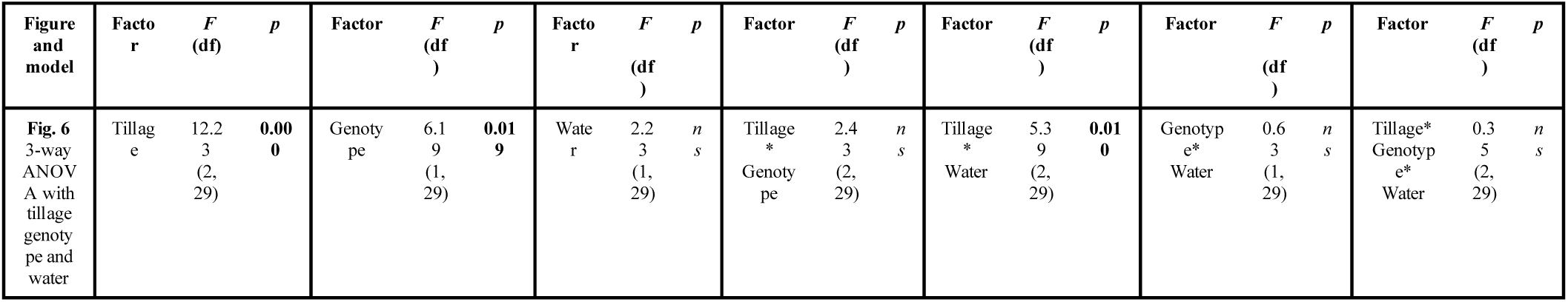
Effects of tillage, genotype, and water treatment on root:shoot ratio after the wheat harvest, analysed by 3-way ANOVA, and corresponding to the mean values presented in Fig. 6. Bold text indicates significant value if *p* <0.05.

Averaging across varieties and water treatments, the direct drilled ley unexpectedly had significantly lower root biomass proportions (0.0795 ± 0.001 s.e.) than both the disc and ploughed (*p* <0.001 in both cases), with no difference between the disc (0.16 ± 0.01 s.e.) and ploughed (0.14 ± 0.002 s.e.) treatments (*p* >0.05). There was no main effect of water treatment, nor was there a significant interaction between wheat variety and tillage, or variety and water treatment, on the root:shoot ratios (*p* >0.05). However, there was a significant interaction between tillage and water treatments (*p* = 0.01) which arose from biomass allocation to roots relative to shoots in the direct drilled ley halving from an arithmetic mean across both wheat genotypes of 0.108 (± 0.009 s.e.) under ambient watering to 0.052 (± 0.008 s.e.), under drought, whereas drought had no effect on the proportion of biomass allocated to roots in the other tillage treatments.

### 3.6 Above and below ground partitioning of ^14^C in wheat three days after radiolabelling

Single wheat tillers taken immediately after exposure to ^14^CO_2_ had high variability of the isotope ranging from 10-99% of the ^14^C supplied, scaling by the numbers of tillers present in each monolith, confirming shoot ^14^C fixation, with no main effects of variety, tillage or water treatments (Supplementary Fig. 3 and Supplementary Table 1). Three days after ^14^C labelling, AxC 169 consistently tended to have a higher percentage of the supplied ^14^C retained in shoots than AxC 157 (Fig. 7a; Table 6), but this was not significant (*p* = 0.077 on log-transformed data). Due to the high variability in the data there were no significant main effects or interaction effects of tillage, water treatment or wheat variety on the percentage of ^14^C supplied recovered in the shoots at this time, which averaged 43.6% across all samples.

**Figure 7.**
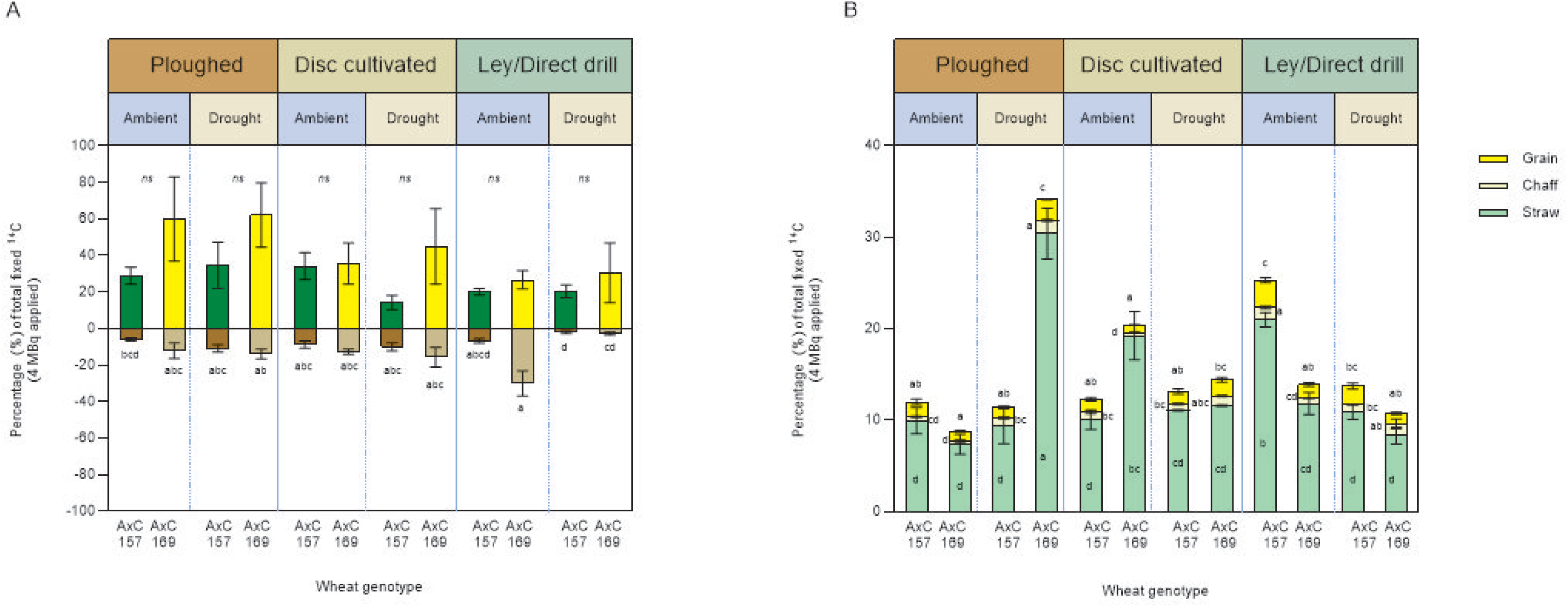
Effects of wheat genotype (AxC 157 & AxC 169), irrigation (control & drought) and tillage (ploughed, disc cultivated, direct drill) on percentage of 4 MBq ^14^C fixed by wheat grown in soil monoliths. A) ^14^C in shoots and roots 3 days after labelling, B) ^14^C in biomass proportions at final harvest (77 days after labelling) Error bars show ± 1 SEM. Means sharing the same letter are not significantly different (*p* > 0.05) (post hoc-Tukey test after 3-way ANOVA.

**Table 6.** Summary of Three-way ANOVA results of main effects of tillage, genotype, and water treatment on wheat mesocosms in ^14^C labelling study for Fig.7a, 7b, and 7c. Bold text indicates significant value if *p* < 0.05

| Figure and model | Plant fraction | Factor | *F* (df) | *p* | Factor | *F* (df) | *p* | Factor | *F* (df) | *p* | Factor | *F* (df) | *p* | Factor | *F* (df) | *p* |  |  |  |
| --- | --- | --- | --- | --- | --- | --- | --- | --- | --- | --- | --- | --- | --- | --- | --- | --- | --- | --- | --- |
| Fig 7a<br>3-way ANOVA with tillage water and genotype (total shoot nat. log) | Total shoot | Tillage | 1.73 (2, 31) | ns | Genotype | 3.34 (1, 31) | ns | Water | 0.45 (1, 31) | ns | Tillage*Genotype | 0.18 (2, 31) | ns | Tillage*Water | 0.40 (2, 31) | ns | Genotype*Water | 0.54 (1, 31) | ns |
|  | Roots | Tillage | 5.005 (2, 28) | **0.013** | Genotype | 12.51 (1, 28) | **<0.001** | Water | 7.91 (1, 28) | ns | Tillage*Genotype | 1.06 (2, 28) | ns | Tillage*Water | 14.81 (2, 28) | **<0.001** | Genotype*Water | 2.38 (1, 28) | ns |
| Fig. 7b<br>3-way ANOVA with tillage genotype and water (grain nat. log) | Grain | Tillage | 5.02 (2, 20) | **0.017** | Genotype | 3.66 (1, 20) | ns | Water | 0.69 (1, 20) | ns | Tillage*Genotype | 5.93 (1, 20) | **0.01** | Tillage*Water | 5.05 (2, 20) | **0.017** | Genotype*Water | 12.93 (1, 20) | **<0.001** |
|  | Chaff | Tillage | 3.07 (2, 21) | ns | Genotype | 0.00 (1, 21) | ns | Water | 18.94 (1, 21) | **<0.001** | Tillage*Genotype | 2.39 (2, 21) | ns | Tillage*Water | 4.78 (2, 21) | **0.019** | Genotype*Water | 27.45 (1, 21) | **<0.001** |
|  | Straw | Tillage | 1.04 (2, 18) | ns | Genotype | 10.12 (1, 18) | **0.005** | Water | 0.19 (1, 18) | ns | Tillage*Genotype | 28.06 (2, 18) | **<0.001** | Tillage*Water | 43.94 (2, 18) | **<0.001** | Genotype*Water | 43.94 (1, 18) | **<0.001** |

In contrast, although ^14^C allocation into roots by day 3 after labelling was generally much lower than retained in shoots (Fig. 7a; Table 6) there were strong genotype effects (ANOVA on natural log transformed data *p* < 0.0001), with AxC 169 putting almost double the percentage ^14^C supplied (10.9%) into roots compared to AxC 157 (6.0%). This was especially the case in the ley monoliths, where averaging across water treatments, roots of AxC 169 received 127% more ^14^C than roots of AxC 157. There was a significant overall effect of tillage (*p* = 0.013), with ^14^C allocation into roots averaging across water treatments, unexpectedly being significantly lower in the ley (5.35%) than in the disc (10.34%) and ploughed treatments (9.57%), which did not significantly differ. There was a significant main effect of water treatment (*p* <0.01), and there were strong interactive effects of tillage with water (*p =* 0.0081) arising from the lack of drought effects on the percentage of supplied ^14^C found in roots in the wheat grown on the ploughed and disc cultivated soil, whereas in the direct drilled ley, drought reduced root allocation from 29% to less than 3% in AxC 169 and from 6.6% to 1.6% in AxC 157.

### 3.7 Allocation of the ^14^C in wheat shoot biomass at harvest

By the harvest time, 77 days after labelling (Fig. 7b) the total ^14^C remaining in shoots averaging across genotypes had decreased by more two thirds to 14.6% compared to the values 3 days after labelling (Fig. 7a). At harvest there were no significant effects of tillage, genotype, and water treatments on total ^14^C in shoot biomass (*p* > 0.05), but there were significant treatment effects on ^14^C in straw, chaff, and grain when analysed separately (Table 6). Straw was the main sink for the remaining shoot ^14^C in both wheat lines, across all treatments, with AxC 169 retaining an average of 14.8% of supplied ^14^C in straw compared to 12.0% in AxC 157, a varietal difference that was highly significant (*p* =0.005), Table 6).

There were no significant main effects of tillage or water treatments, but both variables had significant interactive effects with variety (tillage *p* < 0.001; water *p* = 0.001), with each other (tillage x water treatment *p* < 0.001), and in the three-way interaction of variety by tillage and water (*p* < 0.001). These effects arose from the retaining of relatively large amounts of ^14^C retained in straw in AxC 169 in the ploughed and droughted treatment (30%), in AxC 157 in the ley with ambient water (21%) and in AxC 169in the disc ambient watering treatment (19%).

Although the wheat was only starting to elongate and was not flowering at the point of ^14^C labelling, at the final harvest the chaff and grain in total held an average of 2.4% of the supplied ^14^C, and this was about 16% of the total supplied ^14^C still found in the shoots.

Chaff contained an average of 0.83% of the ^14^C supplied to the plants, with no main effect of wheat variety or tillage treatments. However there was a highly significant effect of the water treatments (*p* < 0.001), and highly significant interaction between variety and water wheat variety on the percentage of ^14^C supplied that was recovered in the chaff, as AxC 157 showed no effect of drought, whereas in AxC 169 drought increased the ^14^C in chaff from 0.41% (± 0.05 s.e.) to 1.28% (± 0.14 s.e.) which were significantly different (*p* < 0.001).

There was also a significant interaction (*p* < 0.019) between the tillage and water treatments, arising from drought increasing the ^14^C in chaff with ploughing, but not with disc cultivation or direct drilling into ley.

At the harvest, ^14^C in grain contained an average of 1.6% of the ^14^C supplied, which was double that in chaff (Fig. 7b). There was no significant effect (Table 6) of variety on the amount of ^14^C found in grain, although there was a trend for this to be higher in AxC 157 than AxC 169 (*p* = 0.070). However, there were significant effects of tillage (*p* < 0.05), and interactions between variety and tillage (*p* = 0.01), variety and water treatment (*p* = 0.01), and between tillage and water treatment (*p* < 0.05). The ^14^C in grain as a percentage of the amount supplied to the plants increased from 1.3% (± 0.14 s.e.) in the disc cultivated to 1.5% (± 0.19 s.e.) in the ploughed to 2.0% (± 0.24 s.e.) in the ley, with the disc and ley treatments being different (*p* < 0.05). The interaction effects of variety and tillage arose from AxC 157 in the ley treatments having higher grain ^14^C content than the ploughed or disc cultivated treatments, but this effect was not seen with AxC 169. An opposing effect of drought on the two wheat varieties gave the variety by water treatment interaction, with grain ^14^C being nearly double under ambient compared to drought conditions in AxC 157, whereas with AxC 169 the ambient condition gave only a quarter of the ^14^C found in the drought treatment grain. The interaction between tillage and water treatment arose from the ley ambient grain having significantly more ^14^C than in the disc cultivated ambient treatment, averaging across varieties.

### 3.8 Residual ^14^C in soil to 15 cm depth at time of harvest

The ^14^C remaining in soil 0-15 cm depth in the monoliths at harvest averaged only 0.65% of the ^14^C supplied to the plants. Except for soil taken from the ambient ley, a greater percentage of ^14^C in soil was found with AxC 169 than AxC 157, irrespective of water treatment (Fig. 8), giving a significant overall varietal (*p* = 0.029) effect (Table 7, Supplementary Table 2). Tillage also had significant effects (*p =* 0.031) with less ^14^C in the ley than both the ploughed and disc cultivated soil. There were no significant effects of water treatment nor any interactions between the variety, tillage, and water treatments on the soil _14C._

**Figure 8.**
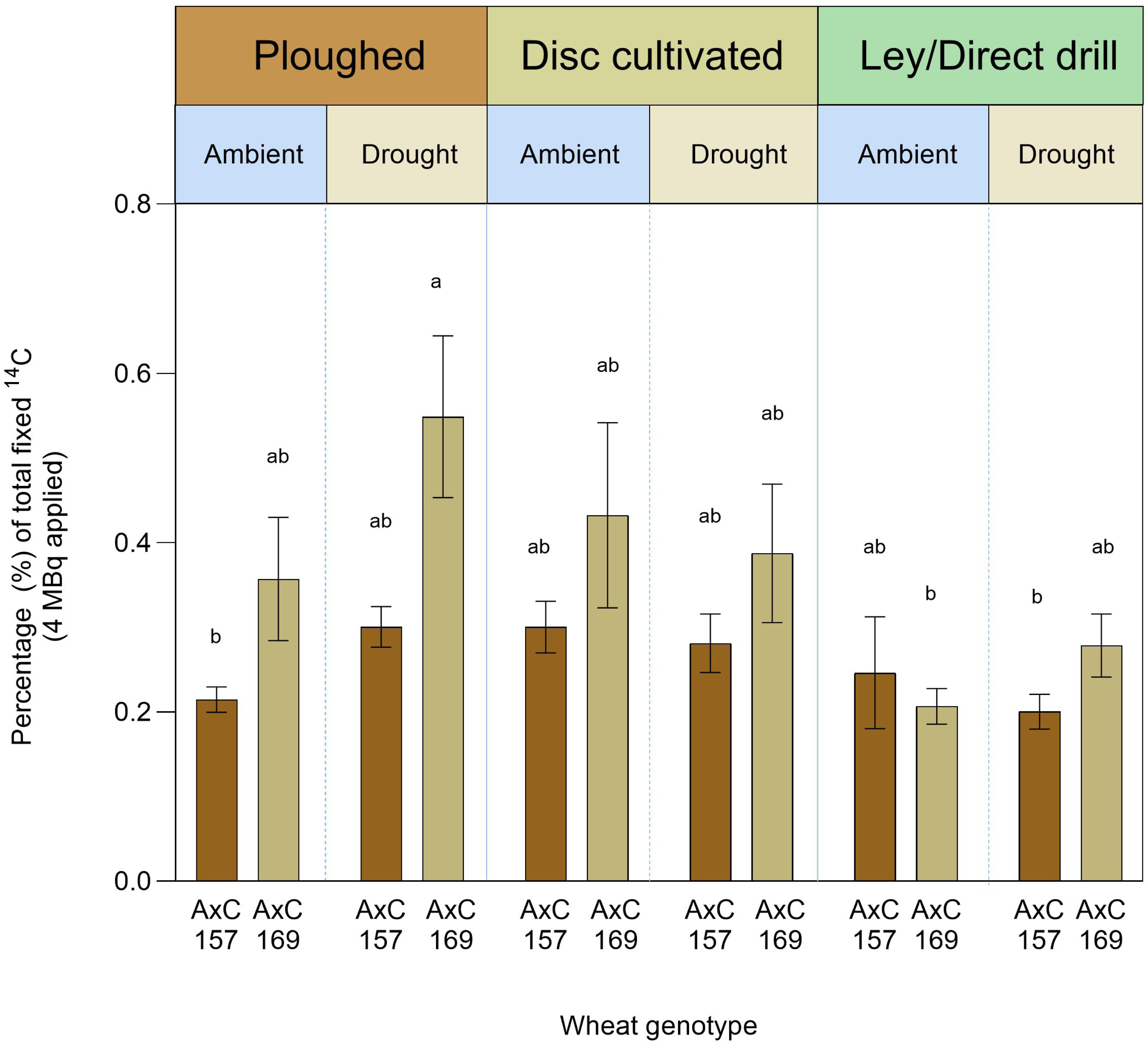
Effects of wheat genotype (AxC 157 & AxC 169), water (control & drought) and tillage (plough, disc cultivated and direct drilled into ley) on percentage of the 4 MBq ^14^C supplied to wheat recovered in soil after the wheat harvest. Error bars show ± 1 SEM. Means sharing the same letter are not significantly different (*p* > 0.05) (post hoc-Tukey test after 3-way ANOVA with tillage, drought and genotype as factors.

**Table 7.**
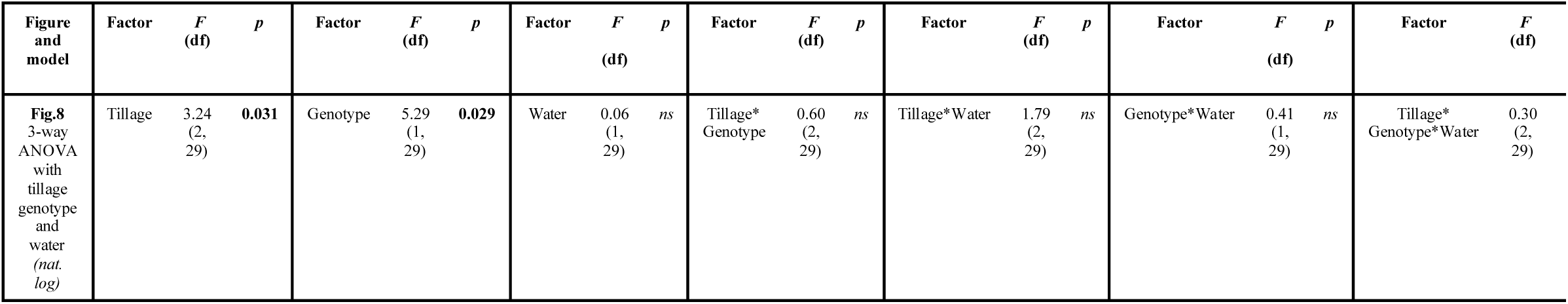
Summary of Three-way ANOVA results of main effects of tillage, genotype, and water treatment on soil from mesocosms in ^14^C labelling study for Fig. 8. Bold text indicates significant value if *p* < 0.05

## 4 Discussion

*4.1 Effects on soil health of wheat genotypes direct drilled into ley compared to wheat grown in permanent arable soil with disc and ploughed cultivation*

Our study corroborates findings of recent research showing the regenerative effects of mown grass-clover leys on soil structure and hydrological functioning (Berdeni *et al.,* 2021, Guest *et al.,* 2022; Hallam *et al.,* 2020) and crop yields (Austen *et al.,* 2022) on the same soil type as the present study. Compared to the ley, reducing tillage intensity from ploughing to disc cultivation for three years in the present study had modest effects on soil structure, with soil compaction assessed by bulk density significantly increasing in the disc cultivated soil 0-7 cm deep (Fig. 3b), but the abundance of > 2 mm WSA slightly increasing (Fig. 4), but this was not significant.

Importantly, we show that direct drilling winter wheat into three-year ley can retain some of the previously reported soil health improvements compared to continuous arable cropping using disc cultivation or ploughing. The former ley soils increased soil water-holding capacity in the early parts of the growing season, reduced bulk density, and substantially increased the proportion of the soil mass in > 2 mm WSA even after growing the winter wheat crop to harvest (Figs. 2-4). However, whilst the two wheat varieties did not differ in their effects on soil water-holding capacity or bulk density, the taller variety (AxC 157), was more effective than the semi-dwarf AxC 169 in retaining the increased > 2 mm WSA from the ley (Fig. 4). Contrary to some previous research (Subira *et. al,* 2016; Velu *et al*., 2017), and our original hypotheses, this effect was not due to greater root biomass produced in the topsoil by the taller variety, since the shorter variety gave the higher root biomass, especially on the ploughed soil, and with no difference between varieties on the ley (Fig. 5).

Gene linkages between wheat height and root traits were investigated in the 199 near-isogenic wheat lines of the Avalon x Cadenza population by Bai *et al.,* (2013) who found some co-located quantitative trait loci, but on screening 25 short (50-75 cm) and 23 tall (120-140 cm) wheat varieties from a more diverse germplasm collection they found shoot height and root proliferation traits were not simply related. Our findings from only two Avalon x Cadenza wheat lines, that differ more modestly in height (24 cm) and fit between the more extreme short and tall categories of Bai *et al.,* (2013), at 85 and 109 cm (Austen *et al.,* 2022), are consistent with their more extensive evidence that wheat dwarfing does not automatically reduce root biomass allocation. In more contrasting wheat genotypes, ranging from 30 cm to 120 cm tall, Gooding *et al.,* (2012) found nitrogen use efficiency followed a quadratic function that peaked in shoots of 80-95 cm tall. When the genotypes were grown organically those with shoots less than 80 cm suffered increasing competition from weeds as stature decreased. Considering the nutrient use efficiency and yields obtained in both conventional and organic management trials, Gooding *et al.,* (2012) saw little benefit in dwarfing cultivar heights below 80 cm, affirming the importance of research focussed on genotypes in the height range we investigated.

The proportion of soil mass in > 2 mm WSA in the ley used to grow AxC 157 under ambient conditions (41%) was almost identical (39%) to that reported by Guest *et al.,* (2022) for three-year grass-clover leys in fields on the same soil type and adjacent to the one used to source the soil monoliths used in the present study, suggesting that direct drilling winter wheat and its growth to harvest the following summer in this case did not result in degradation of the soil structure. This equivalence between studies is also seen in the ploughed arable controls in which > 2 mm WSA comprised 3-6% of the soil mass in the present study and were 7% of the soil mass in Guest *et al.,* (2022). In contrast, when AxC 169 was grown on the ley the abundance of > 2 mm WSA fell to similar values to those seen in the permanent arable disc and ploughed soils without leys in the rotation (Fig. 4). Furthermore, in the droughted ley treatment the biomass of roots of both wheat varieties substantially decreased, but only with AxC 157 was this associated with a decrease in > 2 mm WSA (Figs. 4-5). Whilst the better soil aggregation under the taller wheat variety partially supported our original hypothesis, the mechanism involved could not be attributed to greater root biomass, contrary to our hypothesis.

These findings raise questions about how the root systems of the two wheat varieties might differ in their capacity to maintain, degrade and regenerate soil macroaggregation. For example, genotype effects could arise either directly through root exudates like polysaccharides that assist aggregation that can differ between cereal genotypes (Galloway *et al.,* 2020), or indirectly through differences in genotype effects on mycorrhization (Thirkell *et al.,* 2022), or earthworm populations (Chassé *et al*., 2019), both of which are important agents of soil aggregation (Li *et al.,* 2021; Piotrowski *et al*., 2004). In our previous field trial AxC 157 supported approximately double the mycorrhizal colonization of AxC 169 (Austen *et al.,* 2022), and although these differences were not statistically significant, their magnitude suggests this could have influenced soil aggregation. Earthworm populations were not assessed in the present study, but experimental manipulation of these populations in soil mesocosms of the same size incubated in the SoilBioHedge experiment in fields at Leeds University farm, showed that they increase the abundance of soil aggregates > 250 μm in this soil type by more than 16% (Hallam *et al.,* 2020). In the monoliths of the arable and 2-year grass-clover ley soils studied by Berdeni *et al*., (2021) after a wheat harvest earthworm numbers ranged from 146 individuals m^−2^ in the arable soil, to 285 individuals m^−2^ in the former ley, with no legacy effects of a drought treatment imposed on a subset of these mesocosms.

Importantly, given the wheat varietal differences in soil macroaggregation seen only in the direct drilled ley treatment in the present study, with the increasing use of leys and reduced tillage in regenerative farming systems, further research is needed to ensure that wheat lines are selected not only to yield well in this system, but also to maintain this core component and indicator of soil health (Guest *et al.,* 2022). Soil macroaggregates > 2 mm play an important role in soil organic C sequestration, protecting micro-aggregate bound organic matter from microbial oxidation, so that increased macroaggregation is associated with increased soil organic C and N storage (Guest, *et al.,* 2022; Puerta *et al.,* 2018; Puget *et al.,* 2020; Yu *et al*., 2015). The high demand for N by a wheat crop is likely to accelerate microbial oxidation of soil organic C and N, and in so doing accelerate the turnover of macroaggregates that are held together by temporary organic binding agents (Tisdall & Oades, 1982). Our study of WSA was focussed on upper topsoil (0-7 cm depth, where aggregate stability is most important with respect to impacts of rainfall and infiltration, but our sampling of root biomass in the monolith did not investigate if the two wheat varieties differed in profile distribution of root biomass. Nonetheless, the low root biomass relative to shoots of wheat (Figs. 5-6), is consistent with previous research on other wheat varieties (Sun *et al.,* 2018) and the low root length densities of wheat (Hodgkinson *et al.,* 2017) indicating that wheat is a poor contributor to soil health. This contrasts with grass-clover leys in which the high rooting densities lead to better topsoil structure (Berdeni *et al*., 2021; Zani *et al*., 2021), facilitated by large increases in earthworm populations (Llanos *et al.,* 2025; Prendergast-Miller *et al.,* 2021), boosted especially by the beneficial effects of white clover (Marley *et al.,* 2024). These effects in turn have been shown to deliver improved hydrological functioning, and reduced soil bulk density, as found in the present study, as well as in other long-term ley studies (Jarvis *et al*., 2017) in comparisons with intensively tilled and cropped soils which often experience slumping, coalescence and compaction (Or *et al.,* 2021).

### 4.2 Effects of leys on wheat yields and resilience to drought stress

The regeneration of soil fertility and structure by the legume-rich leys in the present study was evidenced by the wheat grain yields of 3.69-3.76 t ha^-1^ that were increased by 1.63-2.07 t ha^-1^ (Fig. 5) compared to the long-term arable soil growing wheat (123% more than disc cultivated and 77% more than ploughed). These were similar to the yield improvements of 2 t ha^-1^ by wheat grown in a two-year grass-clover ley compared to arable soil from adjacent fields at the same farm reported by Berdeni *et al.,* (2021) using monoliths of the same size as in the present study. The yields achieved in these mesocosm studies using only 35 kg N ha^-1^ fertilizer are approximately 50% of those reported by Austen *et al.,* (2022) for the field trial using the same fertilizer additions on the same plots of ley, which almost reached UK average conventionally produced wheat yields in the study year. They are above the global average of 3.5 t ha^-1^ (Erenstein *et al.,* 2022), but lower than the 4.0-4.5 t ha^-1^ yields obtained from organically grown wheat in the UK (Gooding *et al.,* 2012). The shallow depth of the mesocosms constrains nutrient and water access compared to the deeper field soil, as discussed by Berdeni *et al.,* (2021), although wheat rooting is strongly concentrated in the top 20 cm of soil and exponentially declines with depth with root length densities typically falling to less than 1 cm cm^-3^ below 20 cm (Hodgkinson *et al.,* 2017; Clarke *et al.,* 2017).

Under the drought treatment the former ley experienced the most rapid and extreme drying despite having the greatest initial water-holding capacity (Fig. 2). This was likely due to the greater shoot biomass production on the ley, as reflected in the final harvest, where it was approximately double that of the ploughed and disc cultivated arable soil (Fig. 5), and this would have increased rates of transpiration. Nonetheless, the drought only caused a minor grain yield penalty that was not significant (*p* > 0.05) in the former ley or long-term arable soil monoliths, but the drought did cause a significant decrease in root biomass of both wheat varieties specifically on the ley, giving a substantial increase in root:shoot mass ratio contrary to our initial hypothesis that wheat on the ley would be more resilient to drought. Soil drying and rewetting is known to stimulate soil respiration, reduce soil aggregation and increase organic nitrogen mineralization and nitrification (Zhang *et al.,* 2022). In the first study to investigate how soil drying and rewetting affects rhizodeposition by wheat, Canarini & Dijkstra (2015) found both rhizodeposition and stabilization of new C in soil were reduced, whilst N mineralization was enhanced. In the present study the halving of root mass at harvest in droughted and rewetted ley compared to ambient watered ley may be partly due to increased availability of N mineralized from the greater organic nitrogen co-stored with carbon in soil macroaggregates in these ley soils compared to the permanent arable soils (Guest *et al.,* 2022), reducing the need to invest in root biomass. Importantly, in the context of more frequent and extreme droughts affecting crops, the plasticity in biomass allocation in response to drought stress and rewetting in the former ley was impressive in maintaining grain yields at 93-94% of the ambient watering treatments, whilst the remaining biomass in roots, straw and chaff decreased by 20-24% across the two wheat varieties grown on ley (Fig. 5).

### 4.3 Wheat ^14^C fixation and allocation in biomass and soil

Our study of the effects of ley and tillage management, wheat genotype, and transient drought stress on photosynthate partitioning following ^14^C pulse-labelling during crop stem elongation stage has provided some new insights that were not predicted by previous research or our hypotheses.

Previous field, mesocosm, and pot studies of ^14^C assimilation and partitioning in wheat, were synthesised by Sun *et al.,* (2018). These studies presented data at different growth stages apportioning the ^14^C remaining in the plant-soil system and not respired. Averaging across the 6-7 earlier studies (Atwell *et al.,* 2002; Gregory & Atwell, 1991; Keith *et al.,* 1986; Palta & Gregory, 1997; Qi & Wang 2008; Swinnen *et al.,* 1995a,b) shows that the proportion of the ^14^C remaining in above and below-ground components of wheat after shoot pulse-labelling changes with plant growth stages. During elongation, 76% is retained in shoots rising to 92% at anthesis and 97% at grain filling, with allocation in roots progressively falling from 24% to 8% and 3%. In the present study, 3 days after ^14^C labelling at stem elongation stage, wheat shoots similarly contained on average 72.4% of the total recovered in plants, with 27.6% in roots, consistent with the established rapid allocation of photosynthate to grass and cereal roots within 48 h (Fang *et al*., 2016; Gregory & Atwell, 1991; Leake *et al*., 2006; Sun *et al*., 2018). In the previous studies collated by Sun *et al.,* (2018) the proportion of the total ^14^C remaining (in both wheat and soil) found in soil is much less than in roots, but very variable between studies, and declines from 7.8% (range 1.5-18%) at stem elongation to 3.0% (range 0.1-9.9%) at anthesis and remains about this value at grain filling (3.6%, with a range of 0.2-11.3%). Since we are missing root ^14^C data for the final harvest, we compare the proportion of ^14^C remaining in soil after extraction of roots to that remaining in both shoots and root-free soil (average 4.4% across all treatments) which compares to an average of 10.2% (range 1.9-26%) at stem elongation falling to 3.2% at anthesis, and 3.7% at grain filling in previous studies (Atwell *et al.,* 2002; Gregory & Atwell, 1991; Keith *et al.,* 1986; Palta & Gregory, 1997; Qi & Wang, 2008; Swinnen *et al.,* 1995a,b).

In interpreting these data expressed as percentage of ^14^C remaining in plants and soil it is important to recognize that following shoot fixation the ^14^C remaining in the plant-soil system decreases over time because of metabolic respiration in shoots and roots of non-structural carbohydrates together with microbial oxidation of root exudates and root cell turnover (Garcia *et al*., 2021; Morgan & Austin, 1983; Sun *et al.,* 2018). We found the average ^14^C in wheat shoots of each monolith declined from 1854 kBq immediately after labelling, to 1119 kBq 3 DAL, and to 639 kBq by harvest 77 DAL. Part of the respiratory losses will be generated by the evident mobilization of ^14^C photosynthate captured into vegetative tissues during stem elongation that was recovered at the final harvest in grain and chaff (Fig. 7), showing the strong conservation of photosynthate allocation to reproduction. We did not measure respiratory fluxes, and the amounts of ^14^C recovered from the different plant and soil components therefore under record the net fluxes of ^14^C. In the case of the soil, for example the low amounts recovered may in part be due to higher rates of microbial degradation and respiration from senescent roots and exudates than occur from the shoots which dry on senescence, as previously reported by Palta & Gregory (1997), who found a high rates of ^13^CO_2_ respiration from wheat roots and soil microorganisms under drought.

We originally hypothesised that improvements in soil structure that were expected to be partially retained under the direct-drilled ley would lead to increased ^14^C retained in soil organic matter by harvest time, and for the taller wheat variety to have larger roots and deliver greater amounts of ^14^C into roots and soil especially in the ley soil. Whilst the direct drilled ley did maintain better hydrological functioning than long-term arable soil, and the taller wheat maintained better soil macroaggregation post ley, contrary to both of our hypotheses, the amount of ^14^C retained in the soil was lowest in the ley, and it was the shorter wheat variety, AxC 169, that allocated more ^14^C into roots and soil.

The finding that less of the ^14^C labelled wheat photosynthate was retained in ley soil than the ploughed or disc cultivated soils has important implications for regenerative farming practices that aim to minimize tillage and use leys to sequester soil C (Jordon *et al.,* 2022).

This outcome was unexpected because the greater macroaggregation in the ley soils was expected to result in greater persistence of ^14^C from roots and mycorrhiza, since macroaggregation facilitates the protection of soil organic carbon in microaggregates within macroaggregates (Liu et al., 2021; Yu *et al.,* 2015), with as much as 90% of soil organic carbon in grassland soils occurring in aggregates (Jastrow, 1996). With the greater proportion of the soil mass in macroaggregates in AxC 157 in the ambient ley, we would have expected to see greater ^14^C retained in soil. However, the significant differences in macroaggregation between the ambient and droughted ley with AxC 157 and between the two wheat varieties provided no clear evidence of soil aggregation controlling the fate of the wheat ^14^C photosynthate released via roots in soil (Fig. 8).

One possible explanation for these unexpected findings is priming of saprotrophic microorganisms through the senescence of the ley roots and foliage when treated with herbicide prior to the sowing of the wheat. The accumulated biomass of roots in the three-year ley may have taken some time to decompose with associated increase in the populations of saprotrophic organisms (Baboo *et al.,* 2013) that may then be able to more quickly metabolise ^14^C released from roots and mycorrhizas of the wheat plants than in the permanent arable soils in which the soil microbial biomass is likely to be less. Further research would be needed to confirm this alternative hypothesis.

The effects of wheat genotype on below-ground ^14^C allocation may be simply attributed to the greater root biomass found with AxC 169 than AxC 157 (Fig. 5), resulting in more ^14^C in the roots of the semi-dwarf variety at 3 DAL (Fig. 7a) and remaining in soil at harvest (Fig. 8). However, the large reduction in root biomass found in both wheat varieties in the droughted ley treatment (Fig. 5), which was reflected in the ^14^C allocation to roots 3 days after labelling (Fig. 7a), was not reflected in the ^14^C remaining in the soil at the harvest 77 DAL (Fig. 8), so a direct link between root biomass and soil ^14^C remaining at the harvest time is not established.

One final consideration is potential effects of mycorrhization and carbon delivered into soil via wheat mycorrhizas. Mycorrhizal hyphae in soil are positively associated with both soil macroaggregation and soil organic carbon sequestration (Wilson *et al.,* 2009). This is due both to hyphal enmeshment of soil particles (Tisdall & Oades, 1982) and persistence of a mycorrhizal fungal synthesised substance until recently referred to as “glomalin” that has been found to be a polysaccharide and renamed glomalose (Alptekin *et al.,* 2025). Although the two wheat lines in our previous field trials had shown somewhat contrasted rates of mycorrhization of their roots, and a trend for higher mycorrhization when direct drilled into former ley than when grown on cultivated permanent arable soil (Austen *et al.,* 2022), we have not investigated whether the varieties differ in the lengths of external mycorrhizal mycelium they support in soil or amounts of glomalose produced. Rillig *et al.,* (2002) showed that different plant species grown in the same soil support different lengths of mycorrhizal hyphae, that the mycorrhizal hyphal lengths correlate with the proportion of soil that is held in WSA of 1-2 mm and with the amount of glomalose (glomalin) found in soil. Schweiger & Jacobsen (2000) showed that root length colonisation by mycorrhiza does not predict external hyphal lengths. The disparities that we found between the very small proportion of ^14^C allocated to roots of transiently droughted wheat plants in the ley not being reflected in treatment effects in ^14^C in the soil at harvest might be the result of mycorrhiza-derived C being more important in the relatively small soil ^14^C pools than root inputs, and mycorrhiza inputs may be less affected by drought than roots (Ficken & Warren, 2019). Previous studies using ^13^C tracer have found that droughted plants can preferentially allocate photosynthate to specific mycorrhizal fungal partners (Forczek *et al.,* 2022).

## 5. Conclusion

Our outdoor mesocosm investigation using intact soil monoliths and ^14^C pulse-labelling has enabled the study of growth and partitioning of photosynthate in semi-dwarf and taller wheat on arable soils from field plots on a gradient of soil disturbance from conventional tillage to regenerative direct drilled leys, and in relation to transient drought stress. Despite soil structural improvements of the ley being partially retained, unexpectedly, this did not increase the proportion of ^14^C retained in roots or soil relative to disc or ploughed permanent arable soil. Under transient drought root biomass decreased specifically in the former ley, but yields were maintained by plasticity in photosynthate and biomass allocation. Although the taller variety maintained better soil aggregation in the former ley, contrary to our expectations, its root systems were smaller than those of the semi-dwarf variety. These findings suggest that despite improvements in soil structure and functioning from regenerative agricultural practices C sequestration under wheat crops is not increased, and contrary to some previous reports, wheat genotypes with reduced height do not always have reduced root biomass.

## Author contributions

**Nichola Austen**: Formal analysis, Investigation, Visualization, Validation, Writing – original draft, Writing – review & editing, **Elizabeth Short**: Formal analysis, Investigation, Writing – review & editing, **Steffi Tille**: Investigation, Writing – review & editing, **Irene Johnson**: Formal analysis, Investigation, Writing – review & editing, **Richard Summers**: Resources, Writing – review & editing, **Duncan Cameron**: Conceptualization, Funding acquisition, Methodology, Writing – review & editing, **Jonathan Leake**: Conceptualization, Funding acquisition, Formal analysis, Investigation, Methodology, Project administration, Resources, Supervision, Validation, Visualisation, Writing – original draft, Writing – review & editing.

## Acknowledgements

The setting up and running of the field experiments was only possible through the generous practical assistance and advice of Mr Peter Burgis of NIAB, Headley Hall. We thank Mr Jake Wild also of NIAB Headley Hall for the operation of the plot combine harvester and arranging the data collection of yields from this. Staff at Leeds University farm, including George Sorensen provided essential support for the research. Dr Kirsty Elliott and Dr Dale Bavastock for the collection, transport, and storage of monoliths.

