## Supplementary material for "Regenerative agriculture effects on biomass, drought resilience and ^14^C-photosynthate allocation in wheat drilled into ley compared to disc or ploughed arable soil": Supp Info

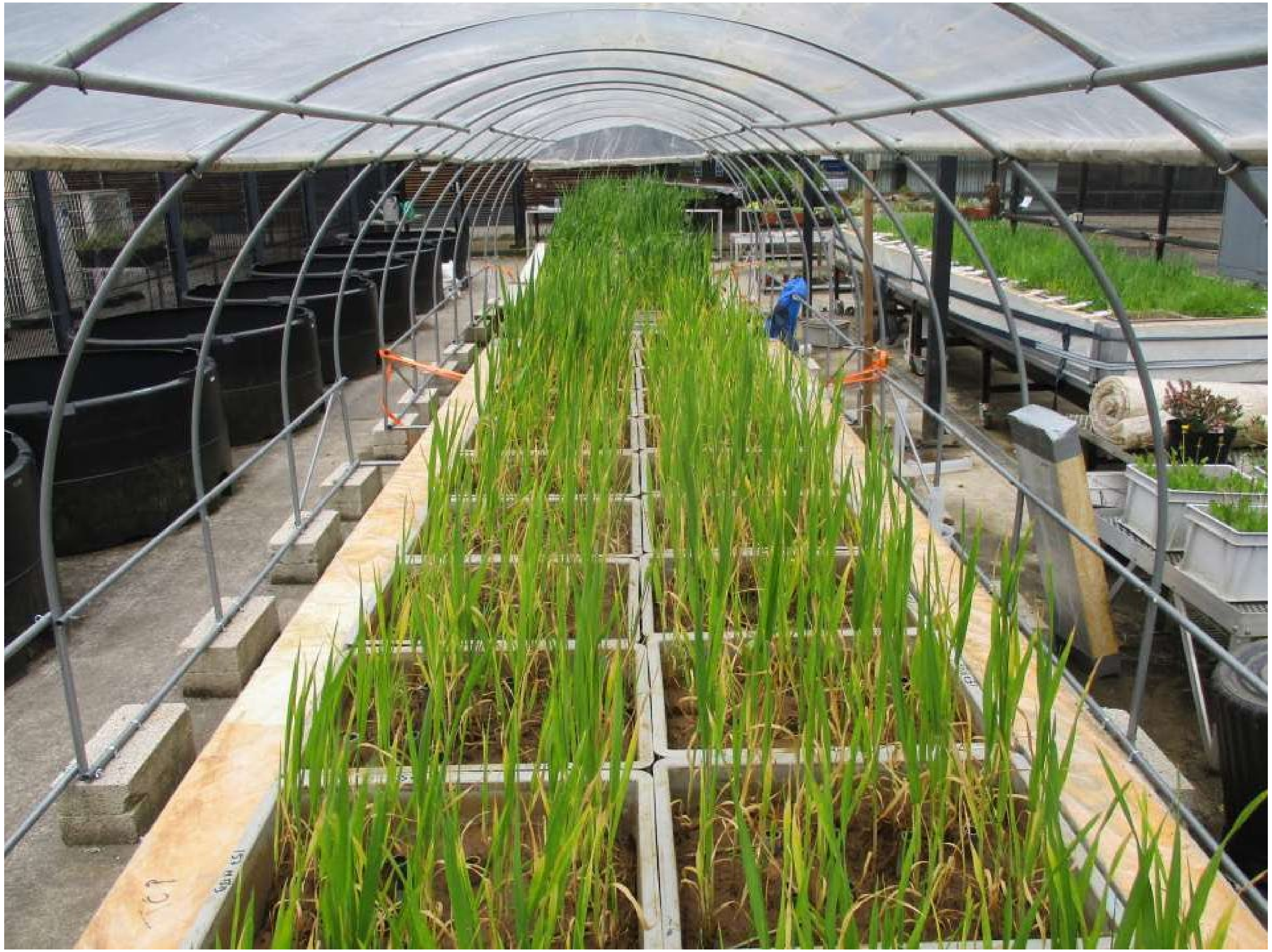

Supp. Figure 1. Monoliths May 2018 immediately after drought treatment. All monoliths brought to field capacity.

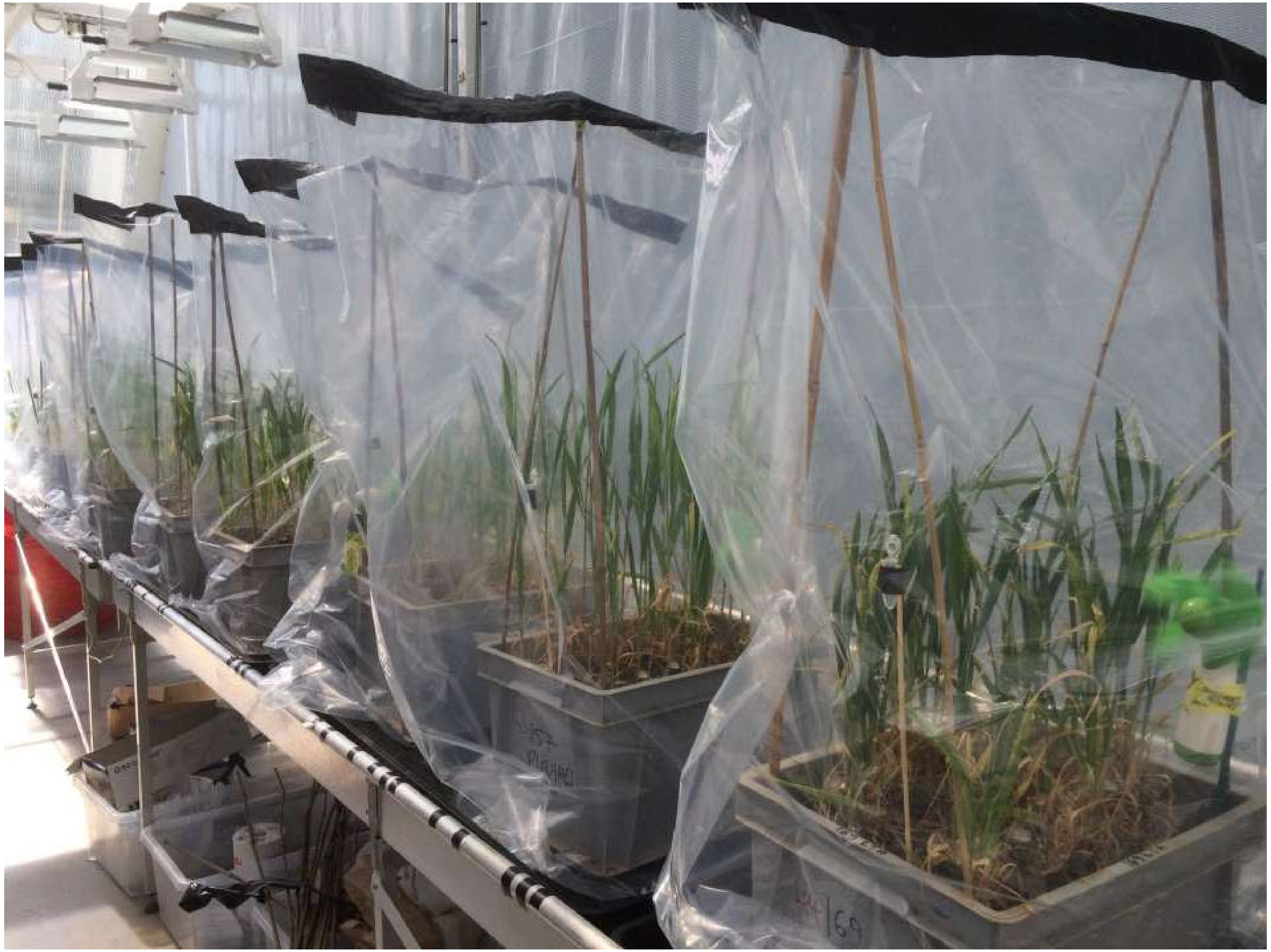

Supp Figure 2. Radiocarbon labelling in May 2018 in a controlled environment greenhouse, with soil monoliths with elongating wheat shoots sealed in clear polythene bags and exposed to 4 MBq $^{14}CO_2$ for 3 hours.

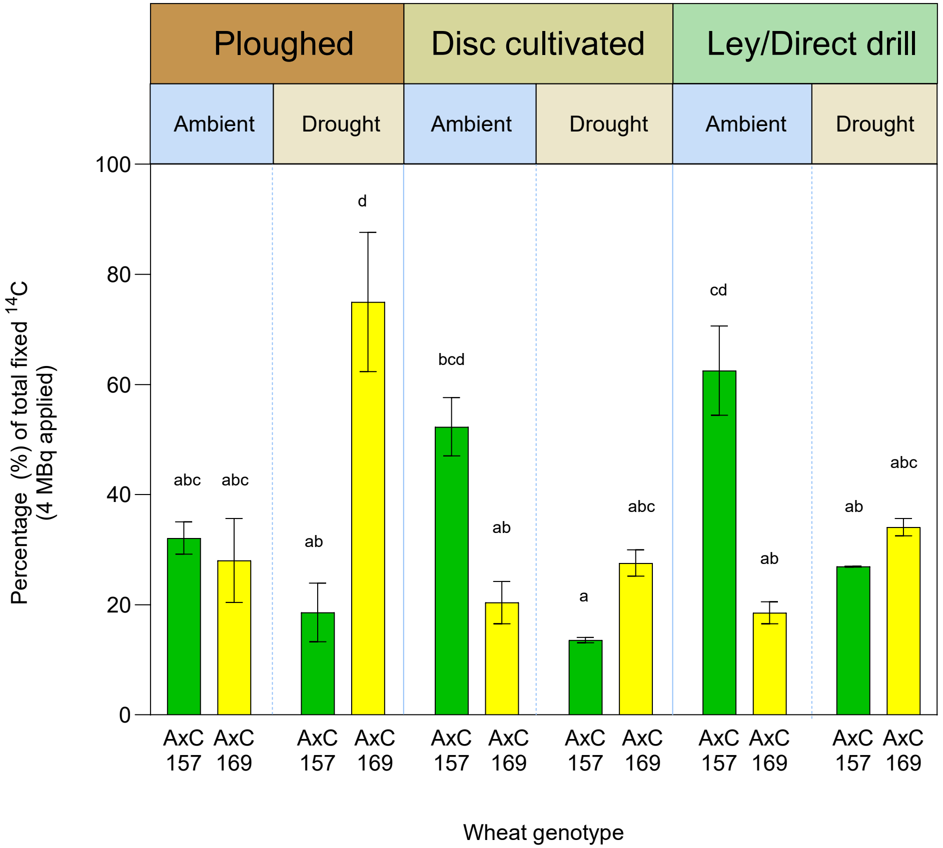


Supp. Figure 3. Effects of wheat genotype (AxC 157 & AxC 169), irrigation (control & drought) and tillage (ploughed, disc cultivated, direct drill) on percentage of 4 MBq $^{14}C$ fixed by wheat grown in soil monoliths. Shoots immediately after labelling

**Supp. Table 1.** Summary of Three-way ANOVA results of main effects of tillage, genotype , and water treatment on wheat mesocosms in 14C labelling study. Bold text indicates significant value if *p = 0.05*

| **Figure and model** | **Plant fraction** | **Factor** | ***F* (df)** | ***p*** | **Factor** | ***F* (df)** | ***p*** | **Factor** | ***F* (df)** | ***p*** | **Factor** | ***F* (df)** | ***p*** | **Factor** | ***F* (df)** | ***p*** | **Factor** | ***F* (df)** | ***p*** | **Factor** | ***F* (df)** | ***p*** |
|---|---|---|---|---|---|---|---|---|---|---|---|---|---|---|---|---|---|---|---|---|---|---|
| **Fig 7a** 3-way ANOVA with tillage water and genotype | Total shoot | Tillage | 2.63 (2, 20) | *ns* | Genotype | 0.01 (1, 20) | *ns* | Water | 0.64 (1, 20) | *ns* | Tillage* Genotype | 12.81 (2, 20) | **0.0155** | Tillage* Water | 7.69 (2, 20) | **0.003** | Genotype* Water | 72.11 (1, 20) | **<0.001** | Tillage* Genotype* Water | 0.35 (2, 20) | *ns* |

**Supplementary Table 2.** Tukey TSD of statistical analyses of paper figures. Three-way ANOVA or mixed-effects model analysis with *p* value reported as significant if <0.05. All results are *post hoc* Tukey tests. Main effects in main text P=ploughed, DC=disc cultivated, DD=direct drilled, a=ambient, d= drought

| Variable/Figure | Analysis | Transform | Treatment | *P* value |
|---|---|---|---|---|
| **2b** | Two-way ANOVA of monolith weight | - | Pa vs. Pd | 0.0261 |
| **3a** | Three-way ANOVA: tillage | - | DD vs. DC<br>DD vs. P | 0.0028<br>0.0034 |
| | Three-way ANOVA: water | | A vs. D | 0.0204 |
| | Three-way ANOVA whole monolith bulk density: tillage*water*genotype | | 157 Dda vs. 157 Dca<br>157 Dda vs. 157 Pa<br>157 Dda vs. 169 DCd | 0.0064<br>0.001<br>0.0055 |
| **3b** | Three-way ANOVA: tillage | - | DD vs. DC<br>DD vs. P | 0.0011<br>0.0056 |
| | Three-way ANOVA 0-7 cm monolith bulk density: tillage*water | | DDd vs. Dca<br>DDd vs. Pa | 0.0032<br>0.462 |
| **4 (>2000 µm)** | Three-way ANOVA: tillage | | DD vs. DC<br>DD vs. P | 0.0004<br>0.0001 |
| | Three-way ANOVA WSA: genotype*tillage | Sqrt | 157 DD vs. 157 DC<br>157 DD vs. 157 P<br>157 DD vs. 169 DD<br>157 DD vs. 169 DD<br>157 DD vs. 169 P | <0.0001<br><0.0001<br>0.0055<br>0.0001<br><0.0001 |
| | Three-way ANOVA WSA: Genotype*tillage*water | | 157 DDa vs. 157 DCa<br>157 DDa vs. 157 DCd<br>157 DDa vs. 157 Pa<br>157 DDa vs. 157 Pd<br>157 DDa vs. 169 DDa<br>157 DDa vs. 169 DDd<br>157 DDa vs. 169 DCa<br>157 DDa vs. 169 DCd<br>157 DDa vs. 169 Pa<br>157 DDa vs. 169 Pd | 0.0001<br>0.0001<br>0.0003<br>0.0000<br>0.0063<br>0.0114<br>0.0019<br>0.0000<br>0.0002<br>0.0000 |
| **4(1000-2000 µm)** | Three-way ANOVA: tillage | | DD vs. DC<br>DD vs. P | 0.0000<br>0.0000 |
| | Three-way ANOVA: water | | A vs. D | 0.0342 |
| | Three-way ANOVA WSA: tillage*water | | Dda vs. DDd<br>Dda vs. DCa<br>Dda vs. DCd<br>Dda vs. Pa<br>Dda vs. Pd<br>DDd vs. DCa<br>DDd vs. DCd<br>DDd vs. Pa<br>DDd vs. Pd | 0.0024<br>0.0000<br>0.0000<br>0.0000<br>0.0000<br>0.0000<br>0.0000<br>0.0000<br>0.0000 |
| | Three-way ANOVA: genotype*water | Sqrt | 169 A vs. 169 D | 0.0008 |
| | Three-way ANOVA: genotype*water*tillage | | 157 DDa vs. 157 DCa<br>157 DDa vs. 157 DCd<br>157 DDa vs. 157 Pa<br>157 DDa vs. 157 Pd<br>157 DDa vs. 169 DDa<br>157 DDa vs. 169 DCa<br>157 DDa vs. 169 DCd<br>157 DDa vs. 169 Pa<br>157 DDa vs. 169 Pd | 0.0001<br>0.0002<br>0.0003<br>0.0000<br>0.0054<br>0.0003<br>0.0001<br>0.0001<br>0.0000 |

**STable 1 cont.** Tukey TSD of statistical analyses of paper figures. Three-way ANOVA or mixed-effects model analysis with *p* value reported as significant if <0.05. All results are *post hoc* Tukey tests. Main effects in main text P=ploughed, DC=disc cultivated, DD=direct drilled, a=ambient, d= drought

| Variable/Figure | Analysis | Transform | Treatment | *P* value |
|---|---|---|---|---|
| **4(1000-2000 μm) Cont.** | Three-way ANOVA: genotype*tillage*water | sqrt | DDd vs. 157 DCa | 0.0000 |
| | | | DDd vs. 157 DCd | 0.0000 |
| | | | DDd vs. 157 PA | 0.0000 |
| | | | DDd vs. 157 Pd | 0.0000 |
| | | | DDd vs. 169 DDa | 0.0391 |
| | | | DDd vs. 169 DCa | 0.0000 |
| | | | DDd vs. 169 DCd | 0.0000 |
| | | | DDd vs. 169 Pa | 0.0000 |
| | | | DDd vs. 169 Pd | 0.0000 |
| | | | 157 Dca vs. 169 DDd | 0.0068 |
| | | | 157 DCd vs. 169 Dda | 0.0000 |
| | | | 157 Pa vs. 169 Dda | 0.0000 |
| | | | 157 Pd vs. 169 Dda | 0.0000 |
| | | | 157 Pd vs. 169 DDd | 0.0016 |
| | | | 169 Dda vs. 169 DDd | 0.0001 |
| | | | 169 Dda vs. 169 DCa | 0.0000 |
| | | | 169 Dda vs. 169 DCd | 0.0000 |
| | | | 169 Dda vs. 169 Pa | 0.0000 |
| | | | 169 Dda vs. 169 Pd | 0.0000 |
| | | | 169 DDd vs. 169 Dca | 0.0359 |
| | | | 169 DDd vs. 169 Pa | 0.0064 |
| | | | 169 DDd vs. Pd | 0.0033 |
| **4 (250-1000 μm)** | Three-way ANOVA: tillage | | DD vs. DC | 0.022 |
| | | | DD vs. P | 0.0006 |
| | | | 157 DD vs. 157 DC | 0.0001 |
| | | | 157 DD vs. 157 P | 0.0000 |
| | Three-way ANOVA: genotype*tillage | | 157 DD vs. 169 DD | 0.0009 |
| | | | 157 DD vs. 169 DC | 0.0003 |
| | | | 157 DD vs. 169 P | 0.0000 |
| | | | 157 Dda vs. 157 DDd | 0.0034 |
| | | - | 157 Dda vs. 157 DCa | 0.0000 |
| | | | 157 Dda vs. 157 DCd | 0.0001 |
| | | | 157 Dda vs. 157 Pa | 0.0000 |
| | Three-way ANOVA: genotype*tillage*water | | 157 Dda vs. 157 Pd | 0.0000 |
| | | | 157 Dda vs. 169 DDa | 0.0003 |
| | | | 157 Dda vs. 169 DDd | 0.0012 |
| | | | 157 Dda vs. 169 DCa | 0.0001 |
| | | | 157 Dda vs. 169 DCd | 0.0001 |
| | | | 157 Dda vs. 169 Pa | 0.0000 |
| | | | 157 Dda vs. 169 Pd | 0.0001 |
| **4 (53-250 μm)** | Three-way ANOVA: tillage | | DD vs. DC | 0.0000 |
| | | | Dd vs. P | 0.0000 |
| | Three-way ANOVA: water | | A vs. D | 0.0005 |
| | | | 157 DD vs. 157 DC | 0.0000 |
| | | | 157 DD vs. 157 P | 0.0000 |
| | Three-way ANOVA: tillage*genotype | | 157 DD vs. 169 DD | 0.0244 |
| | | | 157 DD vs. 169 DC | 0.0000 |
| | | | 157 DD vs. 169 P | 0.0000 |
| | | - | | |
| | | | 157 DDa vs. 157 DDd | 0.0100 |
| | | | 157 DDa vs. 157 Dca | 0.0000 |
| | | | 157 DDa vs.157 DCd | 0.0000 |
| | | | 157 DDa vs.157 Pa | 0.0002 |
| | | | 157 DDa vs.157 Pd | 0.0000 |
| | Three-way ANOVA tillage*genotype*water | | 157 DDa vs.169 DDd | 0.0001 |
| | | | 157 DDa vs.169 DCa | 0.0002 |
| | | | 157 DDa vs.169 DCd | 0.0000 |
| | | | 157 DDa vs.169 Pa | 0.0001 |
| | | | 157 Dda vs. 169 Pd | 0.0001 |
| | | | 157DDd vs. 157 Pd | 0.0271 |

**STable 1 cont.** Tukey TSD of statistical analyses of paper figures. Three-way ANOVA or mixed-effects model analysis with *p* value reported as significant if <0.05. All results are *post hoc* Tukey tests. Main effects in main text P=ploughed, DC=disc cultivated, DD=direct drilled, a=ambient, d= drought

| Variable/Figure | Analysis | Transform | Treatment | *P* value |
|---|---|---|---|---|
| **5 (grain)** | Three-way ANOVA: tillage | - | DC vs. DD | 0.0000 |
| | | | DC vs. P | 0.0428 |
| | | | DD vs. P | 0.0000 |
| | Three-way ANOVA: tillage*genotype*water | | 157 DCa vs. 157 DCd | 0.0001 |
| | | | 157 DCa vs. 157 DDd | 0.0011 |
| | | | 157 DCa vs. 169 DDa | 0.0001 |
| | | | 157 DCa vs. 169 DDd | 0.0004 |
| | | | 157 DCd vs. 157 DDa | 0.0001 |
| | | | 157 DCd vs. 157 DDd | 0.0002 |
| | | | 157 DCd vs. 169 DDa | 0.0000 |
| | | | 157 DCd vs. 169 DDd | 0.0001 |
| | | | 157 DDa vs. 157 Pa | 0.0011 |
| | | | 157 DDa vs. 157 Pd | 0.0013 |
| | | | 157 DDa vs. 169 DCa | 0.0001 |
| | | | 157 DDa vs. 169 DCd | 0.0006 |
| | | | 157 DDa vs. 169 Pa | 0.0116 |
| | | | 157 DDa vs. 169 Pd | 0.0033 |
| | | | 157 DDd vs. 157 Pa | 0.0098 |
| | | | 157 DDd vs. 157 Pd | 0.0109 |
| | | | 157 DDd vs. 169 DCa | 0.0011 |
| | | | 157 DDd vs. 169 DCd | 0.0043 |
| | | | 157 DDd vs. 169 Pd | 0.0263 |
| | | | 157 Pa vs. 169 DDa | 0.0007 |
| | | | 157 Pa vs. 169 DDd | 0.0037 |
| | | | 157 Pd vs. 169 Dda | 0.0008 |
| | | | 157 Pd vs. 169 DDd | 0.0042 |
| | | | 169 DCa vs. 169 DCd | 0.0001 |
| | | | 169 Dca vs. 169 DDd | 0.0004 |
| | | | 169 DCd vs. 169 DDa | 0.0003 |
| | | | 169 DCd vs. 169 DDd | 0.0016 |
| | | | 169 DDa vs. 169 Pa | 0.0071 |
| | | | 169 DDa vs. 169 Pd | 0.0020 |
| | | | 169 DDd vs. 169 Pa | 0.0348 |
| | | | 169 DDa vs. 169 Pd | 0.0105 |
| **5 (chaff)** | Three-way ANOVA: tillage | - | DC vs. DD | 0.0016 |
| | | | DD vs. P | 0.0024 |
| **5 (straw)** | Three-way ANOVA: genotype | - | 157 Vs. 169 | 0.0133 |
| | Three-way ANOVA: tillage | | DC vs. DD | 0.0000 |
| | | | DD vs. P | 0.0000 |
| | Three-way ANOVA: water | | A vs. D | 0.0010 |
| | Three-way ANOVA: genotype*tillage*water | | 157 DCa vs. 157 DDa | 0.0000 |
| | | | 157 Dca vs. 169 Dda | 0.0031 |
| | | | 157 DCd vs. 157 DDa | 0.0000 |
| | | | 157 DCd vs. 157 DDd | 0.0092 |
| | | | 157 DCd vs. 169 DDa | 0.0003 |
| | | | 157 DCd vs. 169 DDd | 0.0356 |
| | | | 157 Dda vs. 157 DDd | 0.0402 |
| | | | 157 Dda vs. 157 Pa | 0.0000 |
| | | | 157 Dda vs. 157 Pd | 0.0000 |
| | | | 157 Dda vs. 169 DCa | 0.0000 |
| | | | 157 Dda vs. 169 DCd | 0.0000 |
| | | | 157 DDa vs. 169 DDd | 0.0115 |
| | | | 157 DDa vs. 169 Pa | 0.0000 |
| | | | 157 DDa vs. 169 Pd | 0.0000 |
| | | | 157 DDd vs. 169 DCa | 0.0029 |
| | | | 157 DDd vs. 169 DCd | 0.0006 |
| | | | 157 DDd vs. 169 Pd | 0..0237 |
| | | | 157 Pa vs/ 169 Dda | 0.0150 |
| | | | 157 Pd vs. 169 Dda | 0.0026 |

**STable 1 cont.** Tukey TSD of statistical analyses of paper figures. Three-way ANOVA or mixed-effects model analysis with *p* value reported as significant if <0.05. All results are *post hoc* Tukey tests. Main effects in main text P=ploughed, DC=disc cultivated, DD=direct drilled, a=ambient, d= drought

| Variable/Figure | Analysis | Transform | Treatment | *P* value |
|---|---|---|---|---|
| **5 (straw) Cont.** | Three-way ANOVA: genotype*tillage*water | - | 169 DCa vs. 169 Dda<br>169 Dca vs. DDd<br>169 DCd vs. 169 DDa<br>169 DCd vs. 169 DDd<br>169 DDa vs. 169 Pa<br>169 DDa vs. 169 Pd | 0.0001<br>0.0120<br>0.0000<br>0.0027<br>0.0059<br>0.0007 |
| **5 (roots)** | Three-way ANOVA: genotype | - | 157 vs. 169 | 0.0067 |
| | Three-way ANOVA: water | | A vs. D | 0.0416 |
| | Three-way ANOVA: genotype*tillage | | 157 DC vs. 169 P<br>157 P vs. 169 P | 0.0124<br>0.0015 |
| | Three-way ANOVA: tillage*water | | DDa vs. DDd | 0.0186 |
| | Three-way ANOVA: tillage*water*genotype | | 157 Pa vs. 169 Pa<br>157 Pa vs. 169 Pd | 0.0211<br>0.0291 |
| **6** | Three-way ANOVA: genotype*tillage*water | Nat. log | 157 Dca vs. 157 DDd<br>157 DDd vs. 169 Dca<br>157DDd vs. 136 DCd<br>157 DDd vs. 136 PA<br>157 DDd vs. 169 Pd<br>169 DDd vs. 169 Pa | 0.0408<br>0.0269<br>0.0262<br>0.0048<br>0.0030<br>0.0200 |
| **7a** | Three-way ANOVA: genotype*water | - | 157 a vs. 157 d<br>157 a vs. 169 a<br>157 d vs. 169 d<br>169 a vs. 169 d | 0.0001<br>0.0003<br>0.0008<br>0.0023 |
| | Three-way ANOVA: tillage*water | | DCd vs. Pd | 0.0067 |
| | Three-way ANOVA: genotype*tillage*water | | 157 Dca vs. 157 DCd<br>157 Dca vs. 157 Pd<br>157 Dca vs. 169 Dca<br>157 DCd vs. 157 Dda<br>157 DCd vs. 169 Pd<br>157 Dda vs. 157 DDd<br>157 Dda vs. 157 Pd<br>157 Dda vs. 169 DCa<br>157 Dda vs. 169 DDa<br>157 Dda vs. 169 Pa<br>157 Pa vs. 169 Pd<br>169 Dca vs. 169 Pd<br>169 DCd vs. 169 Pd<br>169 Dda vs. 169 Pd<br>169 Ddd vs. 169 Pd<br>169 Pa vs. 169 Pd | 0.0086<br>0.0297<br>0.0459<br>0.0007<br>0.0001<br>0.0466<br>0.0003<br>0.0037<br>0.0074<br>0.0247<br>0.0030<br>0.0002<br>0.0034<br>0.0005<br>0.0147<br>0.0011 |

| Variable/Figure | Analysis | Transform | Treatment | *P* value |
|---|---|---|---|---|
| **7b (roots)** | Three-way ANOVA: tillage | Nat. log | DC vs. DD | 0.0136 |
| | | | DD vs. P | 0.0340 |
| | Three-way ANOVA: tillage*water | | Dca vs. DDd | 0.0002 |
| | | | DCd vs. DDd | 0.0001 |
| | | | Dda vs. DDd | 0.0001 |
| | | | DDd vs. Pa | 0.0018 |
| | | | DDd vs. Pd | 0.0001 |
| | Three-way ANOVA: tillage*water*genotype | | 157 Dca vs. 157 DDd | 0.0188 |
| | | | 157 DCd vs. 157 DDd | 0.0061 |
| | | | 157 DDd vs. 157 Pd | 0.0033 |
| | | | 157 DDd vs. 169 DCa | 0.0009 |
| | | | 157 DDd vs. 169 DCd | 0.0008 |
| | | | 157 DDd vs. 169 DDa | 0.0001 |
| | | | 157 DDd vs. 169 Pa | 0.0049 |
| | | | 157 DDd vs. 169 Pd | 0.0007 |
| | | | 169 Dda vs. 169 DDd | 0.0032 |
| | | | 169 DDd vs. 169 Pd | 0.0466 |
| **7c (grain)** | Three-way ANOVA: tillage | Nat. log | DC vs. DD | 0.0132 |
| | Three-way ANOVA: genotype*tillage | | 157 DC vs. 157 DD | 0.0080 |
| | | | 157 DD vs. 157 P | 0.0117 |
| | | | 157 DD vs. 169 Dc | 0.0017 |
| | | | 157 DD vs. 169 DD | 0.0109 |
| | Three-way ANOVA: genotype*water | | 157a vs. 169 a | 0.0033 |
| | | | 169a vs. 169 d | 0.0314 |
| | Three-way ANOVA: tillage*water | | Dca vs. Dda | 0.0028 |
| | Three-way ANOVA: tillage*water*genotype | | 157 Dda vs. 157 Pd | 0.0251 |
| | | | 157 Dda vs. 169 Dca | 0.0004 |
| | | | 157 Dda vs. 169 DDd | 0.0268 |
| | | | 157 Dda vs. 169 Pd | 0.0146 |
| | | | 157 DDd vs. 169 Dca | 0.0207 |
| | | | 169 Dca vs. 169 Pd | 0.0128 |
| **7c (chaff)** | Three-way ANOVA: genotype*water | - | 157 A vs. 169 A | 0.0033 |
| | | | 157 A vs. 169 D | 0.0236 |
| | | | 157 D vs. 169 A | 0.0309 |
| | | | 157 D vs. 169 D | 0.0114 |
| | | | 169 A vs. 169 D | 0.0000 |
| | Three-way ANOVA: tillage*water | | Dca vs. Pd | 0.0080 |
| | | | Dda vs. Pa | 0.0104 |
| | | | DDd vs Pa | 0.0063 |
| | | | Pa vs. Pd | 0.0004 |
| | Three-way ANOVA: tillage*water*genotype | | 157 Dda vs. 157 Pa | 0.0039 |
| | | | 157 Dda vs. 169 Dca | 0.0024 |
| | | | 157 Dda vs. 169 Dda | 0.0198 |
| | | | 157 Dda vs. 169 Pa | 0.0038 |
| | | | 157 Pa vs. 169 DDd | 0.0316 |
| | | | 157 Pa vs. 169 Pd | 0.0016 |
| | | | 169 Dca vs. 169 DDd | 0.0177 |
| | | | 169 Dca vs. 169 Pd | 0.0011 |
| | | | 169 Dda vs. 169 Pd | 0.0086 |
| | | | 169 DDd vs. 169 Pa | 0.0261 |
| | | | 169 PA vs. 169 Pd | 0.0016 |

**STable 1 cont.** Tukey TSD of statistical analyses of paper figures. Three-way ANOVA or mixed-effects model analysis with *p* value reported as significant if <0.05. All results are *post hoc* Tukey tests. Main effects in main text P=ploughed, DC=disc cultivated, DD=direct drilled, a=ambient, d= drought

| Variable/Figure | Analysis | Transform | Treatment | *P* value |
|---|---|---|---|---|
| **7c (straw)** | Three-way ANOVA: genotype*tillage | - | 157 DC vs. 157 DD | 0.0213 |
| | | | 157 DC vs, 169 P | 0.0003 |
| | | | 157 DD vs. 157 P | 0.0037 |
| | | | 157 P vs. 169 DC | 0.0124 |
| | | | 157 P vs. 169 P | 0.0001 |
| | | | 169 DC vs. 169 DD | 0.0307 |
| | | | 169 DD vs. 169 P | 0.0002 |
| | Three-way ANOVA: genotype*water | | 157 D vs. 169 D | 0.0005 |
| | | | 169 A vs. 169 D | 0.0260 |
| | Three-way ANOVA: tillage* water | | Dca vs. DDd | 0.0324 |
| | | | Dca vs. Pa | 0.0054 |
| | | | Dca vs. Pd | 0.0246 |
| | | | DCd vs. Dda | 0.0435 |
| | | | DCd vs. Pd | 0.0004 |
| | | | Dda vs. DDd | 0.0027 |
| | | | Dda vs. Pa | 0.0004 |
| | | | DDd vs. Pd | 0.0000 |
| | | | Pa vs. Pd | 0.0000 |
| | Three-way ANOVA: tillage*water*genotype | | 157 Dca vs. 157 Dda | 0.0026 |
| | | | 157 Dca vs. 169 Dca | 0.0139 |
| | | | 157 Dca vs. 169 Pd | 0.0000 |
| | | | 157 DCd vs. 157 Dda | 0.0164 |
| | | | 157 DCd vs. 169 Pd | 0.0000 |
| | | | 157 Dda vs. 157 DDd | 0.0059 |
| | | | 157 Dda vs. 157 Pa | 0.0024 |
| | | | 157 Dda vs. 157 Pd | 0.0014 |
| | | | 157 Dda vs. 169 DCd | 0.0252 |
| | | | 157 Dda vs. 169 DDa | 0.0147 |
| | | | 157 Dda vs. 169 DDd | 0.0014 |
| | | | 157 Dda vs. 169 Pa | 0.0002 |
| | | | 157 Dda vs.169 Pd | 0.0244 |
| | | | 157 DDd vs. 169 Dca | 0.0306 |
| | | | 157 DDd vs. 169 Pd | 0.0000 |
| | | | 157 Pa vs. 169 Dca | 0.0125 |
| | | | 157 Pa vs/ 169 Pd | 0.0000 |
| | | | 157 Pd vs/ 169 Dca | 0.0074 |
| | | | 157 Pd vs. 169 Pd | 0.0000 |
| | | | 169 Dca vs. 169 DDd | 0.0065 |
| | | | 169 Dca vs. 169 Pa | 0.0011 |
| | | | 169 Dca vs. Pd | 0.0054 |
| | | | 169 DCd vs. 169 Pd | 0.0000 |
| | | | 169 Dda vs. 169 Pd | 0.0000 |
| | | | 169 DDd vs. 169 Pd | 0.0000 |
| | | | 169 Pa vs. 169 Pd | 0.0000 |
| **8** | Three-way ANOVA: tillage | Nat. log | DC vs. DD | 0.0467 |
| | | | DD vs. P | 0.0478 |
